# Optogenetic manipulation of second messengers in neurons and cardiomyocytes with microbial rhodopsins and adenylyl cyclase

**DOI:** 10.1101/2022.10.25.513731

**Authors:** Hanako Hagio, Shiori Hosaka, Wataru Koyama, Aysenur Deniz Song, Janchiv Narantsatsral, Koji Matsuda, Takashi Shimizu, Shoko Hososhima, Satoshi P Tsunoda, Hideaki Kandori, Masahiko Hibi

## Abstract

Even though microbial photosensitive proteins have been used for optogenetics, their use should be optimized to precisely control second messengers *in vivo*. We exploited *GtCCR4* and *KnChR*, cation channelrhodopsins from algae, *BeGC1*, a guanylyl cyclase rhodopsin from a fungus, and photoactivated adenylyl cyclases (PACs) from cyanobacteria (*Oα*PAC) or bacteria (*b*PAC), to control cell functions in zebrafish. Optical activation of *Gt*CCR4 and *Kn*ChR in the hindbrain reticulospinal V2a neurons, which are involved in locomotion, immediately induced swimming behavior, whereas activation of *Be*GC1 or PACs was achieved at a short latency. *Kn*ChR had the highest locomotioninducing activity of all the channelrhodopsins examined. Activation of *Gt*CCR4 and *Kn*ChR in cardiomyocytes induced cardiac arrest, whereas activation of *b*PAC gradually induced bradycardia. *Kn*ChR activation led to an increase in intracellular Ca^2+^ in the heart, suggesting that depolarization caused cardiac arrest. These data suggest that these optogenetic tools can be used to reveal the roles of second messengers in various cell types in vertebrates.

**Impact statement:** We identified efficient and useful microbial channelrhodopsin, guanylyl cyclase rhodopsin, and photoactivated adenylyl cyclase that regulate neural activity and cardiac function in zebrafish.

**Major subject areas:** Neuroscience, Cell biology

**Research organism:** Zebrafish (*Danio rerio*)

## Introduction

Cells perform their functions by responding to various signals. For example, in the nervous system, neurons respond to neurotransmitters to increase or decrease ions and/or chemical mediators such as Ca^2+^, cAMP, and GMP in the cytoplasm (hereafter referred to as second messengers). Similarly, cardiomyocyte function is regulated by sympathetic and parasympathetic nerves that involve noradrenergic and cholinergic receptors, respectively, and control second messengers such as cAMP. To understand the roles of second messengers in cell and tissue functions, it is necessary to manipulate the second messengers at a precise timing and locations, and to examine their effects on cell and tissue functions *in vivo*.

Optogenetics is a rapidly expanding technology that controls or detects cellular functions by using photoreactive proteins that are genetically expressed in cells. Microbial rhodopsins have been used for optogenetics (*Kandori, 2020, 2021*). Microbial rhodopsins bind to chromophore *all-trans* retinal, an aldehyde of vitamin A that is present in most, if not all, animal cells. Upon light absorption, microbial rhodopsins exhibit retinal isomerization from all-*trans* to 13-*cis* retinal and are accordingly converted to active retinal, which is the photoproduct. 13-*cis* retinal is thermally isomerized into all*trans* retinal, and the spontaneous return to the inactive form is termed the “photocycle”. Thus, the activity of microbial rhodopsins can be controlled in the presence or absence of light stimulation. Two main types of microbial rhodopsins are used in optogenetics. The first includes microbial rhodopsins with ion-transporting properties such as light-gated ion channels and light-driven ion pumps, while the other includes microbial rhodopsins with enzymatic activity, i.e., enzymorhodopsins.

Among the ion-transporting rhodopsins, channelrhodopsin 1 and 2 (*Cr*ChR1 and *Cr*ChR2), which are light-gated cation channels, were identified from the green alga *Chlamydomonas reinhardtii (Nagel et al., 2002, Sineshchekov et al., 2002, Suzuki et al., 2003*). When *Cr*ChR1 and *Cr*ChR2 were expressed in *Xenopus* oocytes, they functioned as light-gated cation-selective channels (*Nagel et al., 2002, Nagel et al., 2003). Cr*ChR2 was used to depolarize mammalian cells in response to light (*Boyden et al., 2005, Ishizuka et al., 2006*). Thereafter, variants of *Cr*ChR2 or chimeric forms of *Cr*ChR1 and *Cr*ChR2 were developed to improve the efficiency of expression and induce higher activity than the original *Cr*ChR1 and *Cr*ChR2 (*Berndt et al., 2011, Deisseroth, 2011, Ernst et al., 2014, Wang et al., 2009*). These efforts have made *Cr*ChRs the most commonly used optogenetic tools to induce depolarization in neurons and to mimic neuronal activation through ion channel-type neurotransmitter receptors. However, there are some limitations when using *Cr*ChRs as optogenetic tools. The ion selectivity of *Cr*ChR2 is much higher for H^+^ than Na^+^ (*Nagel et al., 2003*). If pH inside and outside the cell differs, *Cr*ChR2 acts as an H^+^ channel rather than an Na^+^ channel. As *Cr*ChR2 is permeable to Ca^2+^ to some extent (*Nagel et al., 2003*), neuronal activation by *Cr*ChR2 leads to both depolarization and activation of the Ca^2+^ pathway, making it difficult to distinguish between the two effects. *Gt*CCR4 is a light-gated cation channel derived from the cryptophyte *Guillardia theta (Govorunova et al., 2016, Yamauchi et al., 2017*). The light sensitivity of *Gt*CCR4 is higher than that of *Cr*ChR2 while the channel open lifetime lies in the same range as that of *Cr*ChR2 when expressed in mammalian neuronal cells.

Since *GtCCR4* conducts almost no H^+^ and no Ca^2+^ under physiological conditions, *Gt*CCR4 is a high Na^+^-selective channelrhodopsin (*Hososhima et al., 2020, Shigemura et al., 2019). Kn*ChR is another cation channelrhodopsin derived from the filamentous terrestrial alga *Klebsormidium nitens (Tashiro et al., 2021*). Truncation of the carboxyterminal of *Kn*ChR prolonged the channel open lifetime by more than 10-fold, providing strong light-induced channel activity (*Tashiro et al., 2021*). These findings imply that *Gt*CCR4 and truncated variants of *Kn*ChR are alternative optogenetic tools that can compensate for the shortcomings of *Cr*ChRs or display stronger photo-inducing activity than *Cr*ChRs (*Hososhima et al., 2020, Tashiro et al., 2021*).

Among the microbial enzymorhodopsins (*Mukherjee et al., 2019, Tsunoda et al., 2021), BeGC1* is a rhodopsin guanylyl cyclase (Rh-GC) derived from the aquatic fungus *Blastocladiella emersonii*, and is responsible for its zoospore phototaxis (*Avelar et al., 2014). Be*GC1 functions as a light-activated guanylyl cyclase. *Be*GC1 shows a rapid light-triggered increase in cGMP when expressed in *Xenopus* oocytes, mammalian cell lines and neurons, and *Caenorhabditis elegans (Gao et al., 2015, Scheib et al., 2015*).Furthermore, when *Be*GC1 was co-expressed with cyclic nucleotide-gated channel (CNG) in neurons, photoactivation of *Be*GC1 depolarized the neurons and evoked behavioral responses in *C. elegans* (*Gao et al., 2015*), suggesting the feasibility of *Be*GC1-mediated optogenetic control of neural functions.

In addition to enzymerhodopsins, photoactivated adenylyl cyclases (PACs) have also been used to regulate intracellular cyclic nucleotides in cells (*Iseki and Park, 2021*). PACs are flavoproteins that catalyze the production of cAMP in response to light stimulation. PACs from the sulfur bacterium *Beggiatoa* sp. (*b*PAC) (*Losi and Gärtner, 2008*) and the cynobacterium *Oscillatoria acuminata* (*Oa*Pac) (*Ohki et al., 2016*) are well characterized. Both *b*Pac and *Oa*Pac have a BLUF (sensor of blue light using the flavin adenine nucleotide) domain and an adenylyl cyclase catalytic domain. When expressed in *E. coli (Ryu et al., 2010), Xenopus* oocytes, rat hippocampus neurons, and adult fruit flies (*Stierl et al., 2011*), *b*PAC acts as a light-dependent adenylyl cyclase. When *b*PAC was expressed in zebrafish interrenal cells, which is the teleost homologue of adrenal gland cells, cortisol increased in a light-dependent manner (*Gutierrez-Triana et al., 2015). b*PAC was also used for light-dependent control of sperm motility in mice (*Jansen et al., 2015*), the release of neurotransmitter in *C. elegans* neurons (*Steuer Costa et al., 2017*), and the control of developmental processes of *Dictyostelium discoideum (Tanwar et al., 2017*). Compared to *b*PAC, *Oa*PAC showed lower minimum photoactivity in the dark and lower maximum photoactivity upon light stimulation when expressed in HEK293 cells (*Ohki et al., 2016*). Nevertheless, *Oa*PAC induced lightdependent axon growth in rat hippocampal neurons (*Ohki et al., 2016*). These experimental findings indicate that *b*PAC and *Oa*PAC are useful optogenetic tools, although their activity in other cell types and animals are unknown.

In this study, we expressed the channelrhodopsins *Gt*CCR4 and *Kn*ChR, enzymorhodopsin *Be*GC1, and photoactivated adenylate cyclases *b*PAC and *Oa*PAC in hindbrain reticulospinal V2a neurons (*Kimura et al., 2013*), which are involved in the induction of swimming behavior, and in cardiomyocytes using the zebrafish Gal4-UAS system (*Asakawa et al., 2008*), and examined their optogenetic activities. Our findings suggest that the optogenetic control using these tools provides a way to regulate second messengers in vertebrates such as zebrafish *in vivo*.

## Results

### Optogenetic activation of cultured neuronal cells by *Gt*CCR4-3.0-EYFP and epitope-tagged *GtCCR4*

To express rhodopsins *in vivo*, fluorescence markers are useful to confirm expression in target cells. There are two methods for marking rhodopsin-expressing cells: expression as a fusion protein with a fluorescent protein, and expression of epitope-tagged rhodopsin and fluorescent protein separately using a viral 2A (P2A) peptide system. We expressed a fusion protein of *Gt*CCR4-3.0-EYFP, which contains the membrane-trafficking signal and the endoplasmic reticulum (ER)-export signal from a Kir2.1 potassium channel (*Gradinaru et al., 2010, Hoque et al., 2016*), or Myc-tagged *Gt*CCR4 (*Gt*CCR4-MT) and TagCFP separately in neuronal ND7/23 cells, which are a hybrid cell line derived from rat neonatal dorsal root ganglia neurons fused with mouse neuroblastoma (*Wood et al., 1990). Cr*ChR2[T159C]-mCherry was used as a positive control (*Berndt et al., 2011*).As reported for cells expressing *Gt*CCR4-EGFP (*Hososhima et al., 2020, Shigemura et al., 2019, Yamauchi et al., 2017*), irradiation of the ND7/23 cells expressing *Gt*CCR4-3.0-EYFP or *Gt*CCR4-MT-P2A-TagCFP with 511 nm light induced a photocurrent, comparable to irradiation of *Cr*ChR2[T159C]-expressing cells with 469 nm light (Figure 1A). Whereas peak photocurrent was significantly higher than steady-state photocurrent in the cells expressing *Cr*ChR2[T159C]-mCherry, peak and steady-state photocurrents were not significantly different in cells expressing *Gt*CCR4-3.0-EYFP and *Gt*CCR4-MT (Figure 1B), suggesting that photoactivation of *Gt*CCR4 from both constructs immediately reached a peak. Channel closing kinetics after shutting-off light (τ_off_) was equally fast for *Gt*CCR4-3.0-EYFP, *Gt*CCR4-MT, and *Cr*ChR2[T159C]-mCherry (Figure 1C), suggesting that high-frequency photostimulation is possible for these channelrhodopsins. Spectrum analysis revealed that *Gt*CCR4-3.0-EYFP responded to light of slightly longer wavelength than *Cr*ChR2[T159C]-mCherry (Figure 1D). Half saturation maximum (EC_50_) of peak and steady-state photocurrents was lower in *Gt*CCR4-3.0-EYFP than *Cr*ChR2[T159C]-mCherry (Figure 1E), suggesting that *Gt*CCR4-3.0-EYFP responded to weaker light stimulation than *Cr*ChR2[T159C]-mCherry. These data indicate that *Gt*CCR4-3.0-EYFP and *Gt*CCR4-MT are highly sensitive and rapidly reacting tools compared to *Cr*ChR2[T159C]-mCherry and can be activated by slightly red-shifted light in cultured cells.

**Figure 1.**
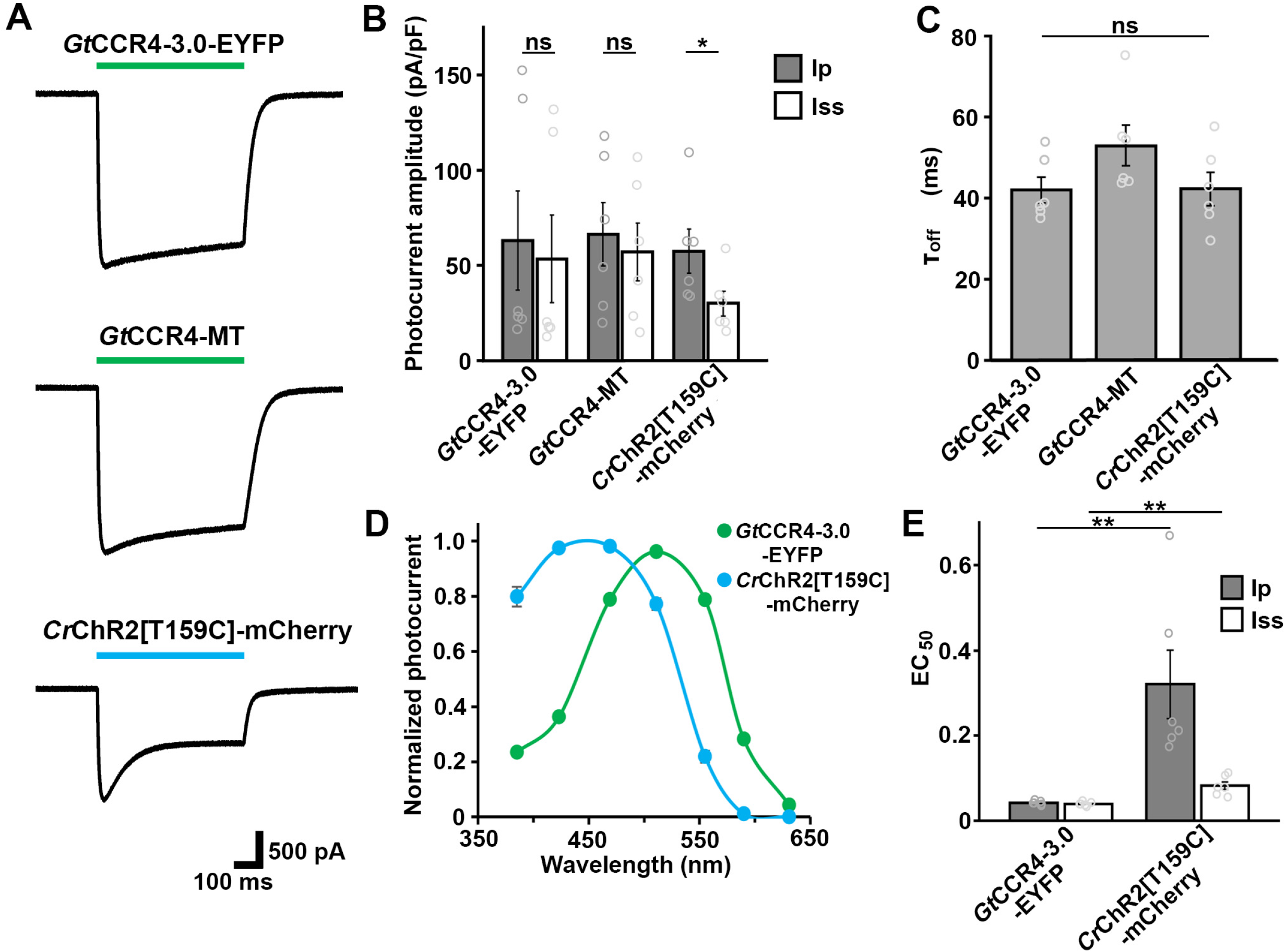
*Gt*CCR4 is capable of depolarizing neuronal cells. (A) Representative photocurrent traces of *Gt*CCR4-3.0-EYFP, *Gt*CCR4-MT-P2A-TagCFP (*Gt*CCR4-MT), and *Cr*ChR2[T159C]-mCherry. Neuronal ND7/23 cells were transfected with the expression plasmids. Electrophysiological recordings were performed. Membrane voltage was clamped at −60 mV. Illumination was with 511 nm light (*Gt*CCR4-3.0-EYFP, *Gt*CCR4-MT) or 469 nm light (*Cr*ChR2[T159C]-mCherry) at 1.4 mW/mm^2^. (B) Photocurrent amplitude. Gray bar: peak photocurrent (Ip); white bar: steady state photocurrent (Iss) (*n*=6, * *p*<0.05). (C) Comparison of the channel closing kinetics after shutting-off light (τ_off_) (*n*=6, * *p*<0.05). (D) Action spectrum of *Gt*CCR4-3.0-EYFP (green circle) and *Cr*ChR2[T159C]-mCherry (Blue circle). Illumination was with 385, 423, 469, 511, 555, 590 or 631 nm at 1.4 mW/mm^2^. Cells were randomly selected for analysis (*n*=5 for *Gt*CCR4-3.0-EYFP, *n*=6 for *Cr*ChR2[T159C]-mCherry). (E) Half saturation maximum (EC_50_) of the peak photocurrent (gray bar) and the steady state photocurrent (white bar) are shown (*n*=6, ** *p*<0.005). Statistical analyses were performed with the Wilcoxon rank sum test. ns, not significant.

### Optogenetic activation of zebrafish locomotion circuit by *GtCCR4* and *KnChR*

In addition to *Gt*CCR4, we also analyzed *in vivo* optogenetic activity of another cation channel *Kn*ChR, whose biophysical properties were previously analyzed using ND7/23 cells (*Tashiro et al., 2021*). We expressed *Gt*CCR4 and *Kn*ChR in the hindbrain reticulospinal V2a neurons in zebrafish, which were reported to drive locomotion (*Kimura et al., 2013*), by using a Gal4-UAS system. As carboxy terminal truncations of *Kn*ChR were shown to prolong the channel open lifetime and give a stronger optogenetic activity (*Tashiro et al., 2021*), a truncated *Kn*ChR containing 272 amino acids from the N-terminus was expressed as a fusion protein containing the membrane-trafficking signal, the ER-export signal, and EYFP (*Kn*ChR-3.0-EYFP). We compared the activities of these channelrhodopsins with that of *Cr*ChR2[T159C]-mCherry. We crossed a transgenic zebrafish *Tg(vsx2:GAL4FF*), which is also known as *Tg(chx10:GAL4*) and expresses a modified transcription factor Gal4-VP16 in the hindbrain reticulospinal V2a neurons (*Kimura et al., 2013*), with Tg lines that express optogenetic tools that are expressed under the control of 5xUAS (upstream activating sequences of yeast *Gal1* gene), the zebrafish *hsp70l* promoter (*Muto et al., 2017*), and mCherry in the heart. Since transgene-mediated protein expression depends on the nature of the introduced gene, the transgene-integrated sites and copy number, we established multiple Tg lines and analyzed stable Tg lines (F_1_ or later generations) with the highest tool expression for each tool. Despite differences in expression between lines, immunohistochemistry with anti-fluorescent protein (anti-GFP and DsRed antibodies for EYFP and RFP/mCherry, respectively) and anti-MT antibodies revealed that in the reticulospinal V2a neurons, *Gt*CCR4-3.0-EYFP, *Kn*ChR-3.0-EYFP, and *Cr*ChR2[T159C]-mCherry were similarly expressed, and GtCCR4-MT was found to be more strongly expressed (Figure 2A, Table 1). We irradiated a hindbrain area of 3-days post fertilization (dpf) Tg larvae expressing *Gt*CCR4, *Kn*ChR, and *Cr*ChR2[T159C] with light of 520, 470, and 470 nm, respectively, for 100 ms (Figure 2B-F, Table 1, Movie 1-4). We measured the rate at which light stimulation induced tail movements (locomotion rate, Figure 2C), the time from stimulation to the onset of tail movements (latency, Figure 2D), the duration of tail movements (Figure 2E), and the amplitude of tail movements (Figure 2F).

**Figure 2.**
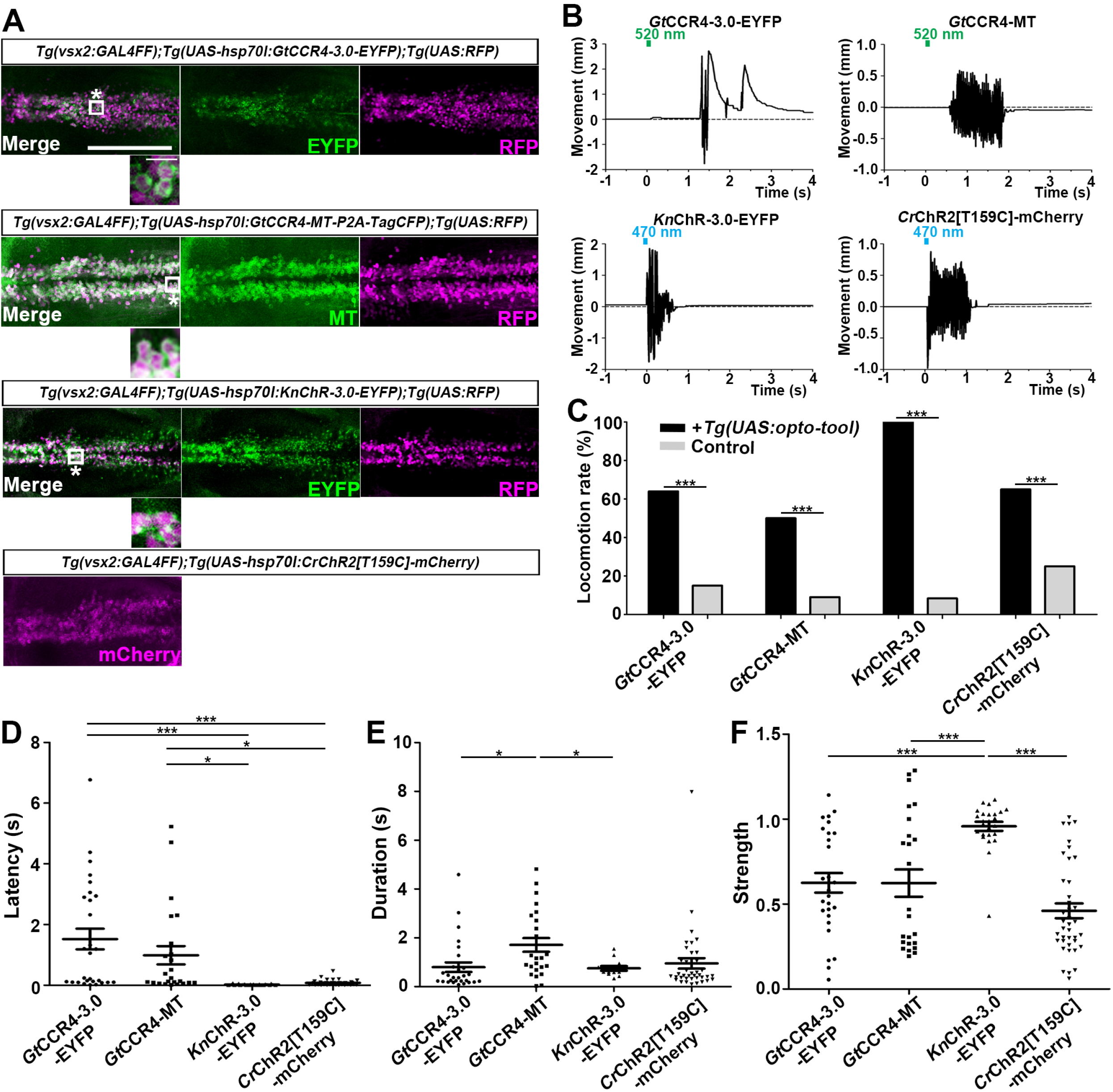
Optogenetic activation of hindbrain reticulospinal V2a neurons by *Gt*CCR4, *Kn*ChR, and *Cr*ChR2[T159C]. (A) Expression of *Gt*CCR4-3.0-EYFP, *Gt*CCR4-MT, *Kn*ChR-3.0-EYFP, and *Cr*ChR2[T159C]-mCherry in the zebrafish hindbrain reticulospinal V2a neurons. 3-dpf (day post fertilization) *Tg(vsx2:GAL4FF);Tg(UAS-hsp70l:GtCCR4-3.0-EYFP*, *GtCCR4-MT-P2A-TagCFP, KnChR-3.0-EYFP*, or *CrChR2[T159C]-mCherry, myl7:mCherry*)larvae were fixed and stained with ant-GFP (EYFP, green), anti-Myc tag (green) or anti-DsRed (RFP, magenta) antibodies. Inset: Higher magnification images for the boxed areas showing double-labeled neurons. In the inset, fluorescence signal intensities were modified to compare the subcellular localization of the tools. (B) Tail movements of 3-dpf Tg larvae expressing *Gt*CCR4-3.0-EYFP, *Gt*CCR4-MT, *Kn*ChR-3.0-EYFP, and *Cr*ChR2[T159C]-mCherry in the reticulospinal V2a neurons after stimulation of the hindbrain area with LED (0.4 mW/mm^2^) light with a wavelength of 520, 520, 470, or 470 nm, respectively, for 100 ms. Light stimulations started at time 0 s. Typical examples are shown. (C) Light stimulation-dependent locomotion rates of 3-dpf Tg larvae expressing *Gt*CCR4-3.0-EYFP, *Gt*CCR4-MT, *Kn*ChR-3.0-EYFP, and *Cr*ChR2[T159C]-mCherry. Six consecutive stimulation trials were analyzed for eight rhodopsin-expressing and nonexpressing (control) larvae of each Tg line. The locomotion rates for *Gt*CCR4-3.0-EYFP, *Gt*CCR4-MT, *Kn*ChR-3.0-EFYP, and *Cr*ChR2[T159C]-mCherry were 63.6% (control 14.6%), 50.0% (control 8.5%), 100.0% (control 8.3%), and 65.0% (control 25.0%), respectively. *** *p* < 0.001, Fisher’s exact test. (D, E, F) Latency (D), duration (E), and strength (F) of light stimulation-evoked tail movements. The time from the start of light irradiation to the first tail movement was defined as latency (s), and the time from the start of the first tail movement to the end of that movement was defined as duration (s). The maximum distance that the caudal fin moved from the midline divided by the body length was measured as its strength. The latency of *Gt*CCR4-3.0-EYFP, *Gt*CCR4-MT, *Kn*ChR-3.0-EFYP, and *Cr*ChR2[T159C]-mCherry was 1.52±0.340, 0.990±0.301, 0.021±0.004, and 0.074±0.028 s, respectively. The duration of *Gt*CCR4-3.0-EYFP, *Gt*CCR4-MT, *Kn*ChR-3.0-EFYP, and *Cr*ChR2[T159C]-mCherry was 0.797±0.198, 1.708±0.282, 0.691±0.054, and 0.950±0.428 s, respectively. The strength of *Gt*CCR4-3.0-EYFP, *Gt*CCR4-MT, *Kn*ChR-3.0-EFYP, and *Cr*ChR2[T159C]-mCherry was 0.625±0.058, 0.623±0.08, 0.958±0.027, and 0.460±0.044 mm, respectively. * *p* < 0.05, ** *p* < 0.01, *** *p* < 0.001, One-way ANOVA with Tukey’s post hoc test. Scale bar = 150 μm in (A), 10 μm in insets of (A).

**Table 1.**
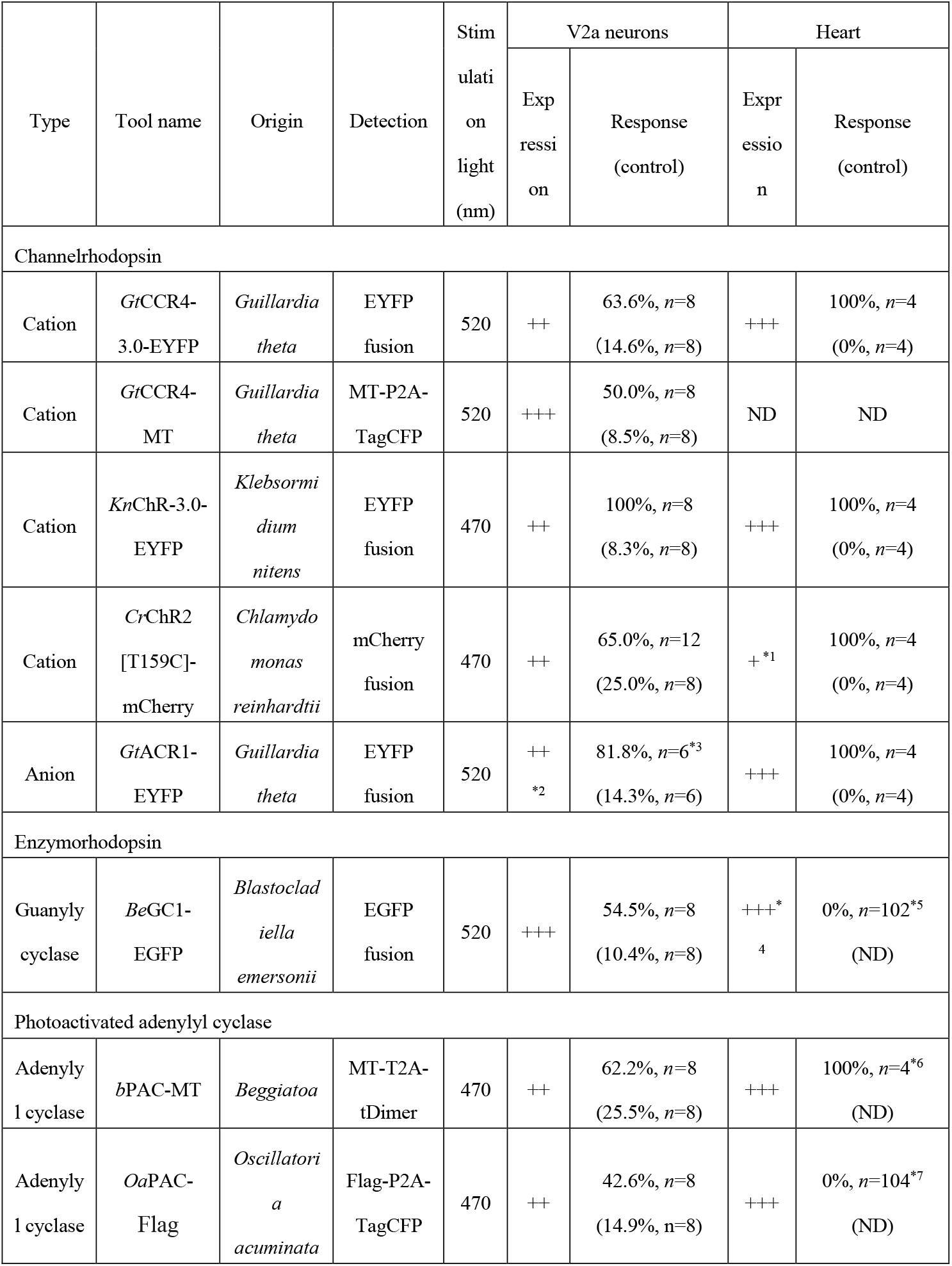
Optogenetic tools. Microbial optogenetic tools were expressed in the hindbrain reticulospinal V2a neurons or cardiomyocytes by the Gal4-UAS system. The expression levels of the tools were determined by immunostaining with anti-tag (MT or Flag) antibodies or anti-fluorescent marker antibodies (anti-GFP and anti-DsRed for EYFP/EGFP and mCherry, respectively) (+ weak, ++ medium, +++ strong expression). The light stimulus-dependent responses (induced swimming or cardiac arrest) are indicated by the percentage of fish that responded to light stimuli. As controls, the responses of sibling larvae that did not express the tools were also examined. *1 Expression of *Cr*ChR2 [T159C]-mCherry was detected by qPCR. *2 Expression was confirmed by detecting EYFP. *3 The percentages of spontaneous tail movements elicited by white light that was inhibited by rhodopsin activation (locomotion-inhibition trials) are indicated (no rhodopsin activation was used as the control). *4 The expression of *Be*GC1-EGFP was determined by observation with an epi-fluorescent microscope MZ16 FA and a fluorescence detection filter (460-500 nm, Leica). *5 Cardiac arrest was not induced with 490-510 nm, 530-560 nm (microscope-equipped light source, *n*=100), or 520 nm (LED) light stimuli (*n*=2). *6 Light stimulation with 470 nm LED light for 5 s induced bradycardia, which took a few minutes to return to normal heartbeats. *7 Stimulation with 460-500 nm (microscope-equipped light source, *n*=100) or 470 nm LED light (*n*=4) induced neither cardiac arrest nor bradycardia, while stimulation with 470 nm LED light occasionally induced transient tachycardia for a few seconds (*n*=2).

Light stimulation of the reticulospinal V2a neurons with *Cr*ChR2[T159C]-mCherry immediately evoked tail movements (Figure 2B, Movie 1). Light stimulation with *Gt*CCR4-3.0-EYFP and *Gt*CCR4-MT evoked tail movements at comparable locomotion rates, although it took more time than *Cr*ChR2[T159C]-mCherry (Figure 2B, C, D, Movie 2, 3). Light stimulation with these channelrhodopsins induced transient tail movements (typically less than 2s, Figure 2E). The strength of the tail movements was not significantly different between *Gt*CCR4-3.0-EYFP/*Gt*CCR4-MT and *Cr*ChR2[T159C]-mCherry (Figure 2F), suggesting comparable photo-inducible activities of *Gt*CCR4 and *Cr*ChR2[T159C] in the reticulospinal V2a neurons. As the activity of *Gt*CCR4-3.0-EYFP was slightly higher than that of *Gt*CCR4-MT (Figure 2C), we used *Gt*CCR4-3.0-EYFP for further analysis. We found that *Kn*ChR-3.0-EYFP was the most potent tool for activating the zebrafish locomotion system among all channelrhodopsins examined. In all trials of all larvae, light stimulation with *Kn*ChR-3.0-EYFP immediately evoked tail movements (Figure 2B, C, D, Movie 4). The strength of evoked tail movements with *Kn*ChR-3.0-EYFP was significantly greater than that with *Gt*CCR4-3.0-EYFP, *Gt*CCR4-MT and *Cr*ChR2[T159C]-mCherry (Figure 2F).

### Optogenetic control of zebrafish heart by *Gt*CCR4 and *KnChR*

We next examined optogenetic activity of *Gt*CCR4 and *Kn*ChR in cardiomyocytes, comparing it with that of the anion channelrhodopsin *Gt*ACR1 (*Gt*ACR1-EYFP), which enables the induction of hyperpolarization in cells (*Govorunova et al., 2015*). We expressed these channelrhodopsins in zebrafish cardiomyocytes by using *Tg(myl7: GAL4FF*), in which GAL4FF was expressed under the promoter of the cardiac myosin light chain gene *myl7*, and the UAS transgenic lines. We established multiple Tg lines for each tool and used Tg lines with the highest tool expression level in cardiomyocytes. Immunostaining with anti-fluorescent protein antibody revealed comparable expression of *Gt*CCR4-3.0-EYFP, *Kn*ChR-3.0-EYFP, and *Gt*ACR1-EYFP in 4-dpf Tg larvae (Figure 3A, Table 1). Stimulation of the entire heart area of 4-dpf Tg larvae with light (520 nm for *Gt*CCR4 and *Gt*ACR1; 470 nm for *Kn*ChR and *Cr*ChR2[T159C]) for 100 ms induced cardiac arrest in all trials, with some differences in latency (Figure 3B, C, Figure 4A, B, Movie 5-8). The latency of cardiac arrest induced by stimulation with these channelrhodopsins was short (less than 1 s), especially that of *Kn*ChR-3.0-EYFP, which was shorter than that of *Gt*CCR4-3.0-EYFP or *Gt*ACR1-EYFP (Figure 4B). Heartbeats (HBs) resumed within 2 s after light stimulation but took longer when stimulated with *Gt*ACR1-EYFP than with *Gt*CCR4-3.0-EFYP or *Kn*ChR-3.0-EYFP (Figure 4C). Considering that *Gt*ACR1 is an anion channelrhodopsin and *Gt*CCR4/*Kn*ChR are cation channelrhodopsins, the mechanisms of cardiac arrest should be different. To address this issue, we monitored intracellular Ca^2+^ concentration in cardiomyocytes using GCaMP6s. Light stimulation with *Kn*ChR-3.0-EYFP increased fluorescence intensity (ΔF/F) of GCaMP6s in the heart and induced cardiac muscle contraction (Figure 4D, Movie 9), whereas that with *Gt*ACR1-EYFP reduced GCaMP6 fluorescence intensity and induced relaxation of the myocardium (Figure 4E, Movie 10), suggesting distinct mechanisms for cardiac arrest induced by the cation channelrhodopsins (*Gt*CCR4 and *Kn*ChR) and anion channelrhodopsin (*Gt*ACR1).

**Figure 3.**
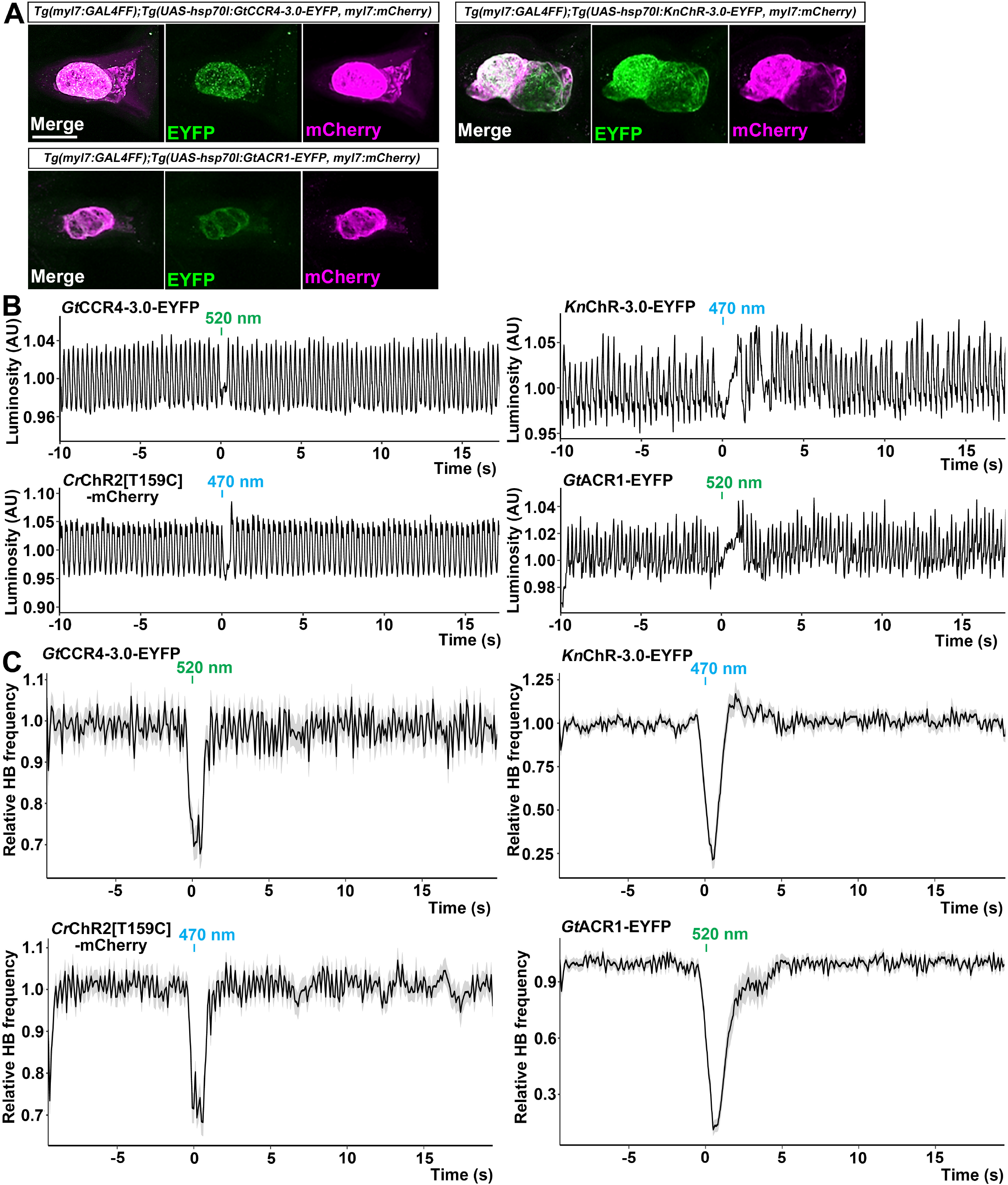
Cardiac arrest induced with *Gt*CCR4-3.0-EYFP, *Kn*ChR-3.0-EYFP, *Cr*ChR2[T159C]-mCherry, and *Gt*ACR1-EFYP. (A) Expression of *Gt*CCR4-3.0-EYFP, *Kn*ChR-EYFP, and *Gt*ACR1-EYFP in cardiomyocytes of *Tg(myl7:GAL4FF);Tg(UAS-hsp70l:GtCCR4-3.0-EYFP*, *KnChR-3.0-EYFP*, or *GtACR1-EYFP, myl7:mCherry*) larvae. 4-dpf Tg larvae were fixed and stained with anti-GFP (for EYFP, green) or anti-DsRed (for mCherry, magenta) antibodies. Z stacks of confocal images. (B) Heartbeat (HB) monitoring by luminosity (AU, arbitrary units) change. The entire heart area of 4-dpf Tg larvae was irradiated with light of appropriate wavelengths (520 nm for *Gt*CCR4 and *Gt*ACR1; 470 nm for *Kn*ChR and *Cr*ChR2) for 100 ms at a strength of 0.5 mW/mm^2^. (C) Average of relative HB frequency. The heart area was irradiated at the indicated periods. Six consecutive stimulus trials were analyzed for four rhodopsin-expressing larvae of each Tg line. Scale bar = 100 μm in (A).

**Figure 4.**
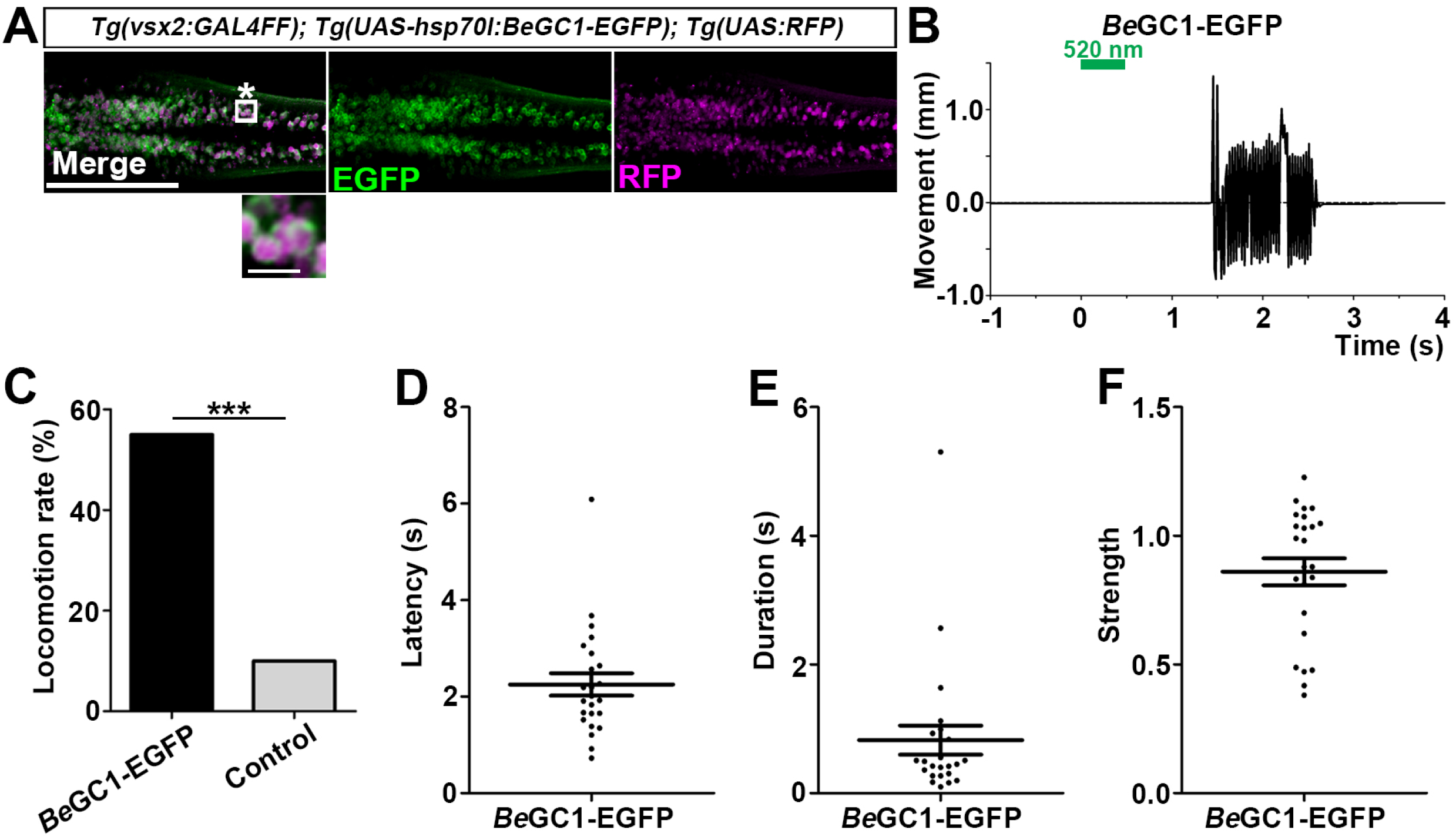
Cardiac arrest and resumption of heartbeats with *Gt*CCR4-3.0-EYFP, *Kn*ChR-3.0-EYFP, *Cr*ChR2[T159C]-mCherry, and *Gt*ACR1-EYFP. (A) Cardiac arrest rates of 4-dpf Tg larvae expressing *Gt*CCR4-3.0-EYFP, *Kn*ChR-3.0-EYFP, *Cr*ChR2[T159C]-mCherry, or *Gt*ACR1-EYFP in cardiomyocytes. The heart area was irradiated with appropriate light (520 nm for *Gt*CCR4 and *Gt*ACR1; 470 nm for *Kn*ChR and *Cr*ChR2) for 100 ms at a strength of 0.5 mW/mm^2^. Sibling larvae that did not express the rhodopsins were used as controls. Six consecutive stimulation trials were analyzed for four rhodopsin-expressing larvae and four control larvae of each Tg line. The cardiac arrest rates for *Gt*CCR4-3.0-EYFP, *Kn*ChR-3.0-EYFP, *Cr*ChR2[T159C]-mCherry, and *Gt*ACR1-EYFP were 100% (control 0%), 100% (control 0%), 100% (control 0%), and 100% (control 0%), respectively. \*\*\**p* < 0.001, Fisher’s exact test. (B, C) Latency to cardiac arrest (B) and time to resumption of HBs (C) after light stimulation with *Gt*CCR4-3.0-EYFP, *Kn*ChR-3.0-EYFP, *Cr*ChR2[T159C]-mCherry, or *Gt*ACR1-EYFP. HB data were obtained from the experiments described above (A). The latencies for *Gt*CCR4-3.0-EYFP, *Kn*ChR-3.0-EYFP, *Cr*ChR2[T159C]-mCherry and *Gt*ACR1-EYFP were 163±15, 55±11, 112±11, and 243±39 ms, respectively. The times to resumption for *Gt*CCR4-3.0-EYFP, *Kn*ChR-3.0-EYFP, *Cr*ChR2[T159C]-mCherry and *Gt*ACR1-EYFP were 418±50, 742±37, 284±19, and 1350±162 ms, respectively. ** *p* < 0.01, *** *p* < 0.001, one-way ANOVA with Tukey’s post hoc test. (D, E) Changes in fluorescence intensity of GCaMP6s (ΔF/F) in the heart of 4-dpf Tg larvae expressing *Kn*ChR-3.0-EYFP and GCaMP6s (D), or *Gt*ACR1-EYFP and GCaMP6s (E). Sibling larvae that did not express the rhodopsins were used as the control. The heart area of Tg larvae were stimulated with a fluorescence detection filter (excitation 470-495 nm, emission 510-550 nm). Two rhodopsin-expressing larvae (green) and two control larvae (black) were analyzed for each rhodopsin. Three trials were analyzed for each larva. *** *p* < 0.001, linear mixed-effects model.

### Optogenetic control of cAMP/cGMP by *BeGC1* and PACs

We examined optogenetic activity of the fungal guanylyl cyclase rhodopsin *Be*GC1 and compared it to that of the flavoprotein-type bacterial photoactivated adenylyl cyclases *b*PAC and *Oa*PAC. By using *Tg(vsx2:GAL4FF*) and UAS Tg lines, we established Tg lines that express *Be*GC1-EGFP, *b*PAC-MT, and *Oa*PAC-Flag in the reticulospinal V2a neurons (Figure 5A, 6A). Although the levels of expression of *b*PAC-MT and *Oa*PAC-Flag were slightly lower than that of *Be*GC1, light stimulation of the reticulospinal V2a neurons with *Be*GC1-EGFP, *b*PAC-MT, and *Oa*PAC-Flag for 500 ms evoked relatively high frequency tail movements, although with a latency of a few seconds (Figure 5B, 5C, 6B, 6C, Movie 11-13). The tail movements induced by activation with *Be*GC1, *b*PAC, or *Oa*PAC had a long latency, but similar duration and intensity compared to activation with the channelrhodopsins (Figure 5D-F, 6D-F). When a cAMP fluorescent indicator Flamindo 2 was co-expressed with *b*PAC-MT or *Oa*PAC-Flag in postmitotic neurons by the *elavl3* promoter and the *elavl3* promote-driven GAL4-VP16 Tg line, continuous light stimulation reduced the ΔF/F of Flamindo 2, which is indicative of an intracellular increase in cAMP (*Odaka et al., 2014*), in the optic tectum (Figure 6G, H). We also used Tg lines expressing *Be*GC1, *b*PAC or *Oa*PAC in cardiomyocytes. Although light stimulation of cardiomyocytes with *Be*GC1-EGFP or *Oa*PAC (Table 1) induced neither cardiac arrest nor bradycardia, activation with *b*PAC for 5 s gradually reduced HBs and it took a few minutes to return to normal HBs (Figure S1, Movie 14). The data indicate that *Be*GC1 and *OaPAC* can be used for optogenetic activation of neurons but not cardiomyocytes, while *b*PAC can be used for optogenetic control of neurons as well as cardiomyocytes in zebrafish.

**Figure 5.**
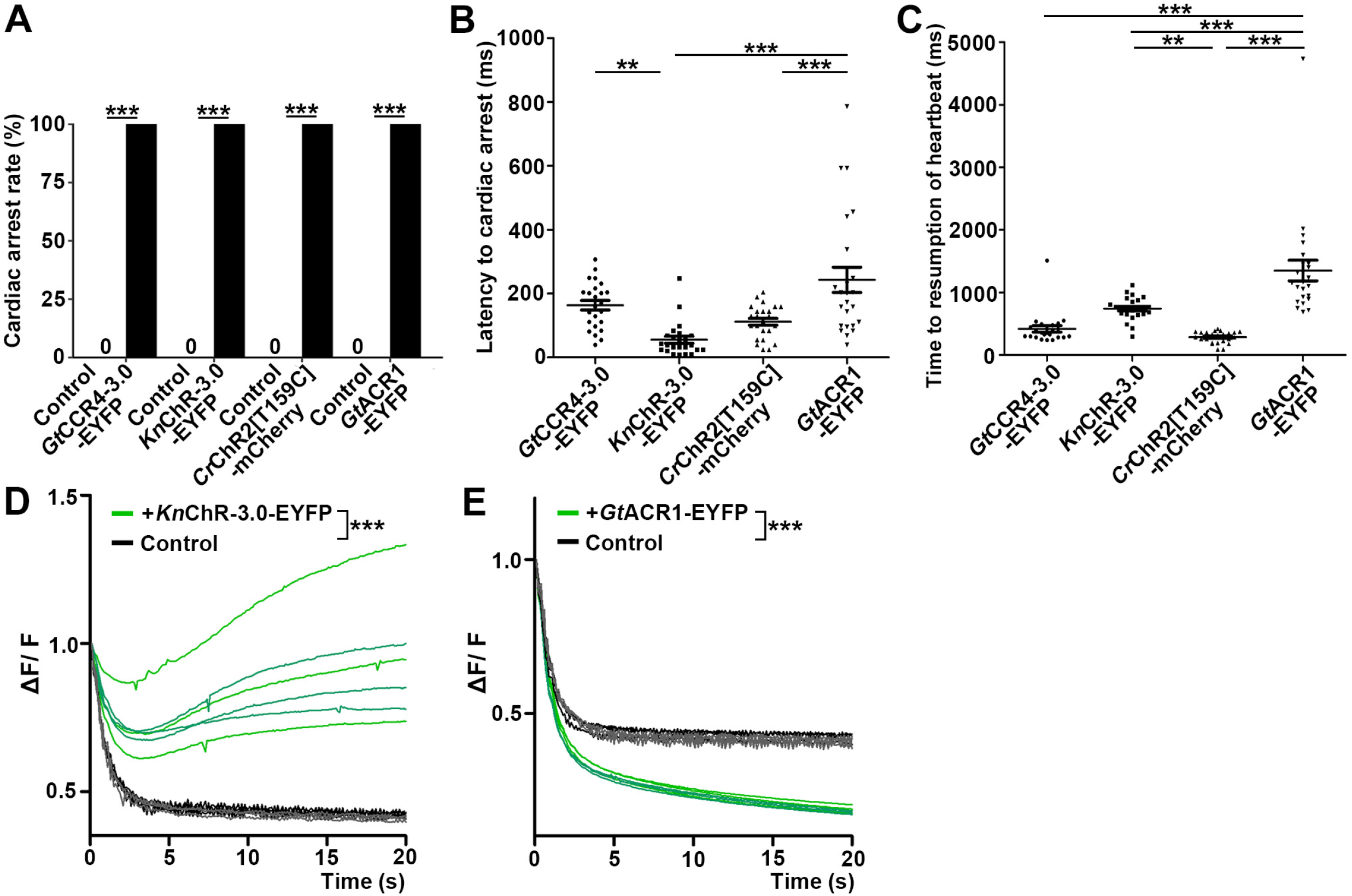
Optogenetic activation of hindbrain reticulospinal V2a neurons with *Be*GC1-EGFP. (A) Expression of *Be*GC1-EGFP in the hindbrain reticulospinal V2a neurons. 3-dpf *Tg(vsx2:GAL4FF);Tg(UAS:BeGC1-EGFP, myl7:mCherry);Tg(UAS:RFP*) larvae were fixed and stained with anti-GFP (EGFP, green) or anti-DsRed (RFP, magenta) antibodies. Inset: higher magnification images for the boxed area showing double-labeled neurons. In the inset, fluorescence signal intensities were modified to compare the subcellular localization of the tools. (B) Tail movements of 3-dpf Tg larvae expressing *Be*GC1 in the reticulospinal V2a neurons after stimulation with light (0.4 mW/mm^2^) at 520 nm for 500 ms. The stimulation started at time 0 s. A typical induced tail movement is shown. (C) Light stimulation-dependent locomotion rates of 3-dpf *Be*GC1-expressing larvae or nonexpressing sibling control larvae. The locomotion rate for *Be*GC1 was 54.5% (control 10.4%). *** *p* < 0.001, Fisher’s exact test. (D, E, F) Latency (D), duration (E), and strength (F) of induced tail movements in the *Be*GC1-expressing larvae. Eight *Be*GC1-expressing larvae (C-F) and eight non-expressing larvae (C) were analyzed. Six consecutive stimulation trials were analyzed for each larva. The latency for *Be*GC1 was 2.25±0.230 s. The duration for *Be*GC1 was 0.825±0.224 s. The strength for *Be*GC1 was 0.860±0.0526 mm. Scale bar = 150 μm in (A), 10 μm in the insets of (A).

**Figure 6.**
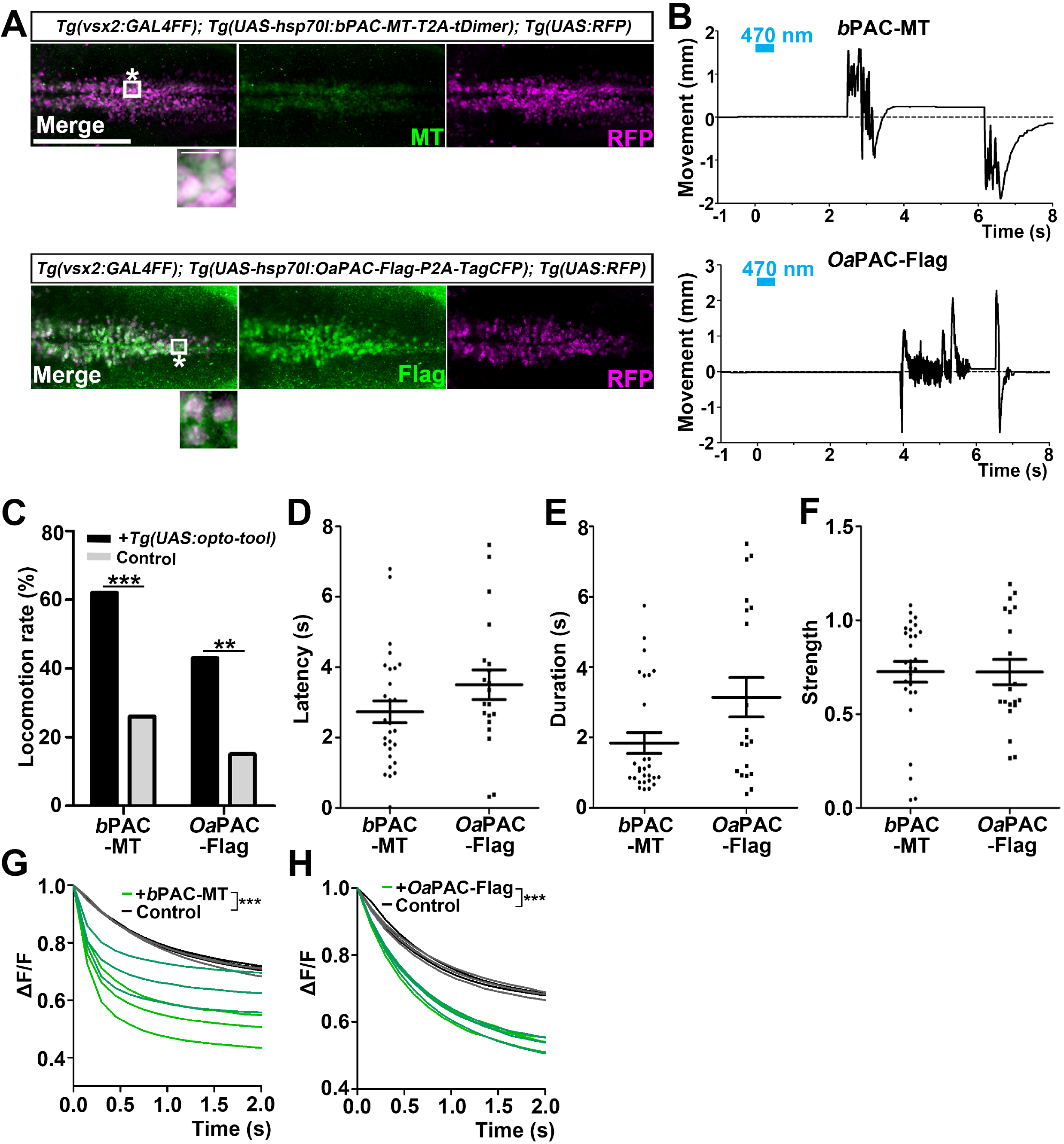
Optogenetic activation of reticulospinal V2a neurons with *b*PAC and *Oa*PAC. (A) Expression of *b*PAC and *Oa*PAC in reticulospinal V2a neurons. 3-dpf *Tg(vsx2:GAL4FF);Tg(UAS-hsp70l:bPAC-MT-T2A-tDimer*, *myl7:mCherry);Tg(UAS;RFP*) and *Tg(vsx2:GAL4FF);Tg(UAS-hsp70l: OaPAC-Flag-P2A-TagCFP, myl7:mCherry);Tg(UAS:RFP*) larvae were fixed and stained with anti-Myc or anti-Flag (green), and anti-DsRed (RFP, magenta) antibodies. To detect relatively weak fluorescent signals of *b*PAC-MT and *Oa*PAC-Flag, images were taken with increased laser power (2x for *b*PAC-MT and 4x for *Oa*PAC-Flag compared to *Be*GC1-EGFP in Figure 5A). Inset: higher magnification images for the boxed areas showing double-labeled neurons. In the inset, fluorescence signal intensities were also modified to compare subcellular localization of the tools. (B) Tail movements of 3-dpf Tg larvae expressing *b*PAC-MT or *Oa*PAC-Flag in the reticulospinal V2a neurons after stimulation with light (0.4 mW/mm^2^) at 470 nm for 500 ms. The stimulation started at time 0 s. Typical examples are shown. (C) Light-induced locomotion rates. Larvae that did not express PACs were used as controls. Six consecutive stimulation trials for eight PAC-expressing and eight control larvae were analyzed. The locomotion rates for *b*PAC and *Oa*PAC were 62.2% (control 25.5%) and 42.6% (control 14.9%). ** *p* < 0.01, *** *p*< 0.001, Fisher’s exact test. (D, E, F) Latency (D), duration (E), and strength (F) for light stimulus-induced tail movements in larvae expressing *b*PAC-MT or *Oa*PAC-Flag. The latency for *b*PAC-MT and *Oa*PAC-Flag was 2.73±0.311 and 3.50±0.422 s, respectively. The duration for *b*PAC-MT and *Oa*PAC-Flag was 1.84±0.297 and 3.14±0.562 s, respectively. The strength for *b*PAC-MT and *Oa*PAC-Flag was 0.726±0.0553 and 0.724±0.0671 mm, respectively. (G, H) Changes in fluorescence intensity (ΔF/F) of cAMP indicator Flamido2 in neurons of Tg larvae expressing *b*PAC-MT (G) or *OaPAC-*Flag (H) after light stimulation. The entire optic tectum area of 3-dpf *Tg(elavl3:GAL4-VP16); Tg(elavl3:Flamindo2); Tg(UAS-hsp70l:bPAC-MT-T2A-tDimer*) and *Tg(elavl3:GAL4-VP16); Tg(elavl3:Flamindo2); Tg(UAS-hsp70l:OaPAC-Flag-P2A-TagCFP*) larvae was stimulated with a fluorescence detection filter (excitation 470-495 nm, emission 510-550 nm). The fluorescence intensity of the optic tectum was measured, and ΔF/F was calculated. Sibling larvae that did not express PACs were used as controls. Three trials for two PAC-expressing (green) and two control (black) larvae were analyzed and the data from a total of six trials were plotted on graphs (\*\*\**p* < 0.001, linear mixed-effects model). Scale bar = 150 μm in (A), 10 μm in insets of (A). Scale bar = 150 μm in (A), 10 μm in insets of (A).

## Discussion

### Utility of channelrhodopsins *Gt*CCR4 and *Kn*ChR

As was reported for *Gt*CCR4-EGFP (*Hososhima et al., 2020, Shigemura et al., 2019, Yamauchi et al., 2017*), *Gt*CCR4-3.0-EYFP and *Gt*CCR4-MT were more sensitive to light stimuli than *Cr*ChR2[T159C]-mCherry in cultured mammalian neuronal cells (Figure 1). However, the optogenetic ability of *Gt*CCR4-3.0-EYFP and *Cr*ChR2[T159C]-mCherry to induce tail movements in the reticulospinal V2a neurons was comparable (Figure 2C, F), and *Gt*CCR4-3.0-EYFP took longer to initiate tail movements (Figure 2D). There are a couple of explanations for this difference. First, the expression of *Gt*CCR4-3.0-EYFP and *Cr*ChR2[T159C]-mCherry proteins might be different. Differences in levels of mRNA expression between Tg lines cannot be ruled out. In addition, there might be difference in translation and cell surface trafficking between these two rhodopsins in zebrafish neurons. *Gt*CCR4-3.0-EYFP contains a membranetrafficking signal and an ER-export signal for expression on the cell surface, but *Gt*CCR4-3.0-EYPF proteins might aggregate slightly within the cytoplasm and might not express efficiently on the cell surface of reticulospinal V2a neurons (Figure 2). Differences in ion channel properties might also affect their activity *in vivo*. *Cr*ChR2 is permeable to not only Na^+^ but also H^+^ and Ca^2+^ whereas *Gt*CCR4 is relatively specific to Na^+^ (*Nagel et al., 2003, Shigemura et al., 2019*). Activation of reticulospinal V2a neurons with *Cr*ChR2 induces an influx of Na^+^, H^+^, and Ca^2+^, while activation with *Gt*CCR4 induces only an influx of Na^+^. This may account for the difference in light-evoked tail movements between *Gt*CCR4 and *Cr*ChR2. On the other hand, the Na^+^-specific channel property of *Gt*CCR4 may favor distinguishing depolarization effects from intracellular Ca^2+^ signaling. Furthermore, the activation wavelength of *Gt*CCR4 is slightly more red-shifted than that of *Cr*ChR2 (Figure 1D) (*Hososhima et al., 2020, Nagel et al., 2003, Shigemura et al., 2019*), which might be useful when used in conjunction with short-wavelength optogenetic tools or neural activation sensors.

We found that *Kn*ChR was the most potent optogenetic tool in zebrafish reticulospinal V2a neurons among the examined tools, which include channelrhodopsins, enzymerhodopsins (in this study), and animal G protein-coupled bistable rhodopsins (in the accompanying paper). Truncation of *Kn*ChR prolongs the channel open lifetime by more than 10-fold (*Tashiro et al., 2021). Kn*ChR conducts various monovalent and bivalent cations, including H^+^, Na^+^, and Ca^2+^, while *Kn*ChR has a higher permeability to Na^+^ and Ca^2+^ and a higher permeability ratio of Ca^2+^ to Na^+^ than *Cr*ChR2 (*Tashiro et al., 2021*). These properties may contribute to the high photo-inducible activity of *Kn*ChR. Activation of *Kn*ChR may induce influx of more cations and depolarization of cells than *Cr*ChR2. Furthermore, since *Kn*ChR can be activated by light with a short wavelength (maximal sensitivity between 430 and 460 nm), *Kn*ChR can be used in conjunction with other red-shifted optogenetic tools and cell activity sensors.

The photoactivation of both cation (*Gt*CCR4, *Kn*ChR) and anion channelrhodopsins (*Gt*ACR1) induced cardiac arrest (Figure 3 and 4). However, activation of *Kn*ChR and *Gt*ACR1 increased and decreased intracellular Ca^2+^ in cardiomyocytes, respectively (Figure 4). Since Ca^2+^ is a readout of depolarization in cardiomyocytes, the data suggest that activation of the cation channelrhodopsins depolarizes cardiomyocytes, increases intracellular Ca^2+^ concentration, and inhibits cardiac resumption, while activation with the anion channel rhodopsins hyperpolarizes cardiomyocytes, decreases intracellular Ca^2+^ concentration, and inhibits cardiac contraction. Tg larvae with a high expression of *Kn*ChR-3.0-EYFP in the heart always showed cardiac arrest after light stimulation (Figure 3, 4), while some larvae with relatively low levels of expression showed accelerated HBs (data not shown). This finding indicates that *Kn*ChR is a strong and tunable optogenetic tool. By altering the degree of depolarization by *Kn*ChR by changing the expression level or the intensity of light stimulation, the function of cardiomyocytes and other cells may be precisely controlled. We also found that light stimulation of hindbrain cholinergic neurons with *Kn*ChR could induce small tail movements (data not shown). Although further studies are needed to elucidate the detailed mechanisms, highly sensitive *Kn*ChR has the potential to identify neural circuits that have not been previously identified with other optogenetic tools.

In this study, we demonstrated that *Gt*CCR4, *Kn*ChR, and *Cr*ChR2 function in both reticulospinal V2 neurons and cardiomyocytes in zebrafish. Given the different ionchannel properties of *Gt*CCR4, *Kn*ChR, and *Cr*ChR2, they can be used for optogenetic manipulation of cell activities in a variety of applications in zebrafish.

### Utility of enzyme rhodopsin *BeGC1* and bacterial flavoprotein PACs

cAMP and cGMP are major second messengers that regulate multiple biological functions in a variety of tissues. We expressed *Be*GC1 and *b*PAC/*Oa*PAC to manipulate intracellular cGMP and cAMP signaling in reticulospinal V2a neurons and cardiomyocytes (Figure 5, 6, S1). Light stimulation of the V2a neuron with *Be*GC1 as well as *b*PAC/*Oa*PAC induced tail movements with a longer delay than with channelrhodopsins. There are two possible mechanisms by which cyclic nucleotides control cell excitability. One mechanism is through cyclic nucleotide-gated ion (CNG) channels, in which binding of cGMP or cAMP to CNG channels opens the cation channels and depolarizes the cell (*Bradley et al., 2005, Matulef and Zagotta, 2003*). The other is through cAMP-dependent protein kinase (PKA), in which binding of cAMP to the regulatory unit of PKA releases the catalytic unit of PKA, resulting in phosphorylation of cation channels such as the voltage-dependent Ca^2+^ channel CaV1.2 (*Fu et al., 2014, McDonald et al., 1994, Reuter, 1983*). The former mechanism is often used for various sensory systems in the nervous system and the latter for the sympathetic noradrenergic regulation of HBs. Which mechanism is used may depend on the availability of necessary components to activate these mechanisms. It is likely that the CNG-mediated mechanism is involved in activation of reticulospinal V2a neurons. In this mechanism, neurons are not activated until the intracellular cGMP concentration reaches the threshold for CNG activation. Consistent with this, activation with *Be*GC1 and PACs induced neural activation with a short delay (Figure 5, 6). On the other hand, the PKA-mediated pathway may be involved in the heart. Activation of *b*PAC but not *Be*GC1 or *Oa*PAC in the heart induced bradycardia (Figure S1). A prolonged increase in intracellular Ca^2+^ induced by PKA-mediated phosphorylation of the Ca^2+^ channel might contribute to the long-lasting bradycardia. Different effects of *b*PAC and *Oa*PAC activation on neurons and the heart are due to differences in basal and photo-inducible activity of these two PACs (*Ohki et al., 2016*). Further analysis is required to reveal the precise mechanism for optogenetic control of cell functions with *Be*GC1 and *b*PAC/*Oa*PAC.

Cell and tissue functions are regulated by various intercellular signaling molecules, such as neurotransmitters, hormones, and cytokines, whose signals are mediated by multiple second messengers. We demonstrated the usefulness of multiple types of channelrhodopsins, cGMP/cAMP-producing tools (in this study), and of GPCR rhodopsins (in the accompanying paper) to manipulate second messenger signaling in zebrafish neurons and cardiomyocytes. Optogenetic studies with these tools alone or in combination will elucidate detailed mechanisms of cellular and tissue regulation through intracellular second messengers.

## Material and methods

### Key resources table

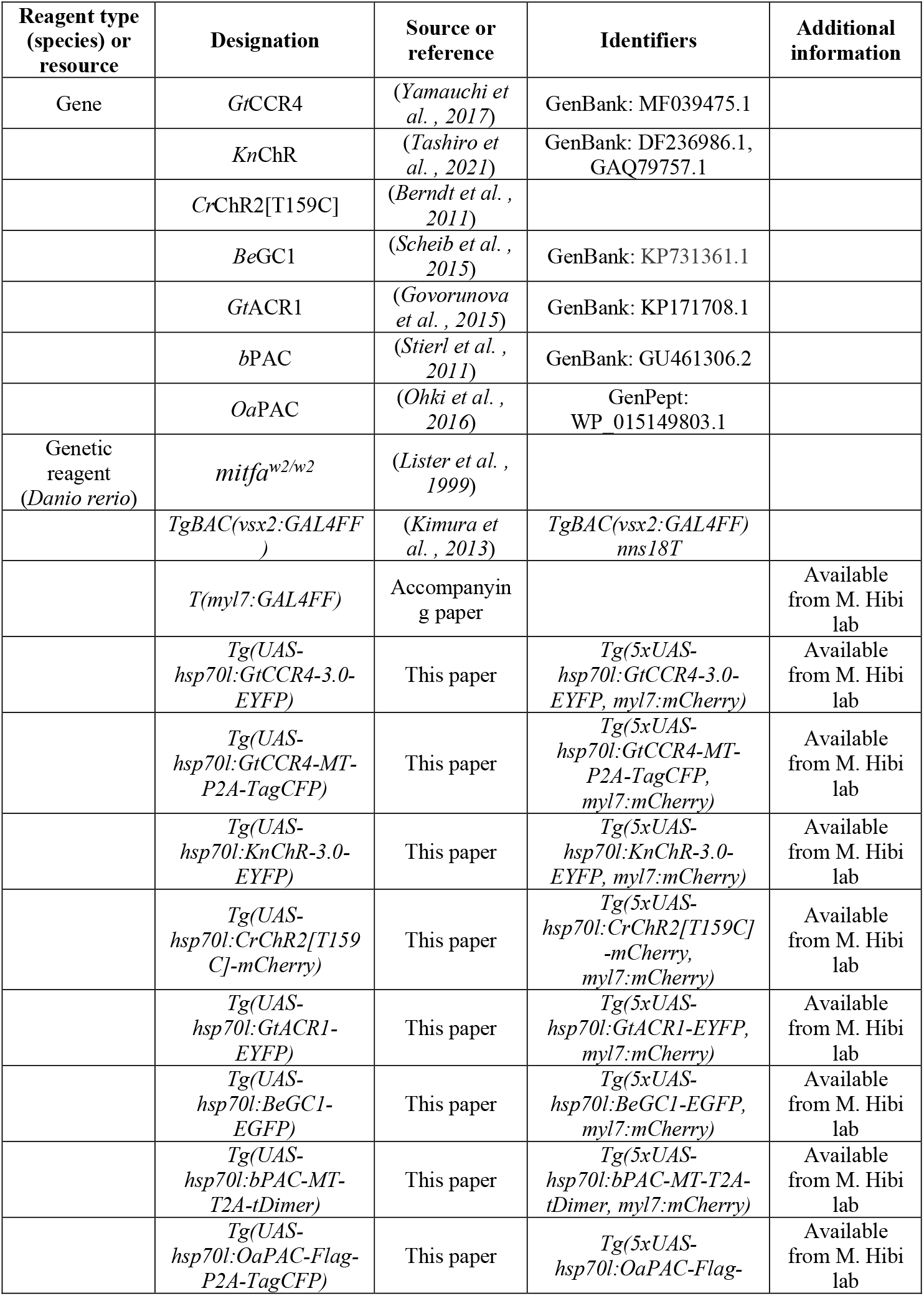
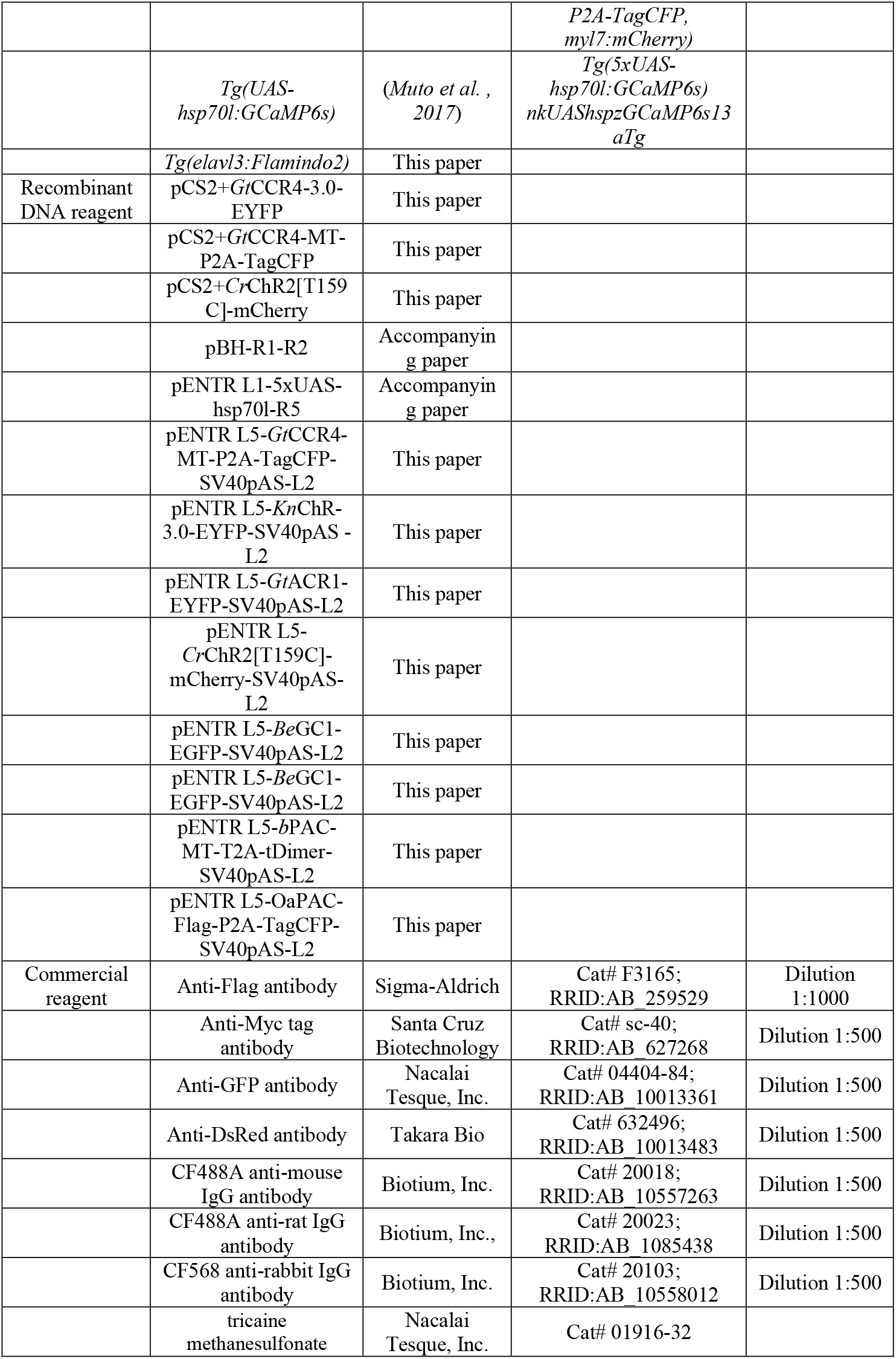
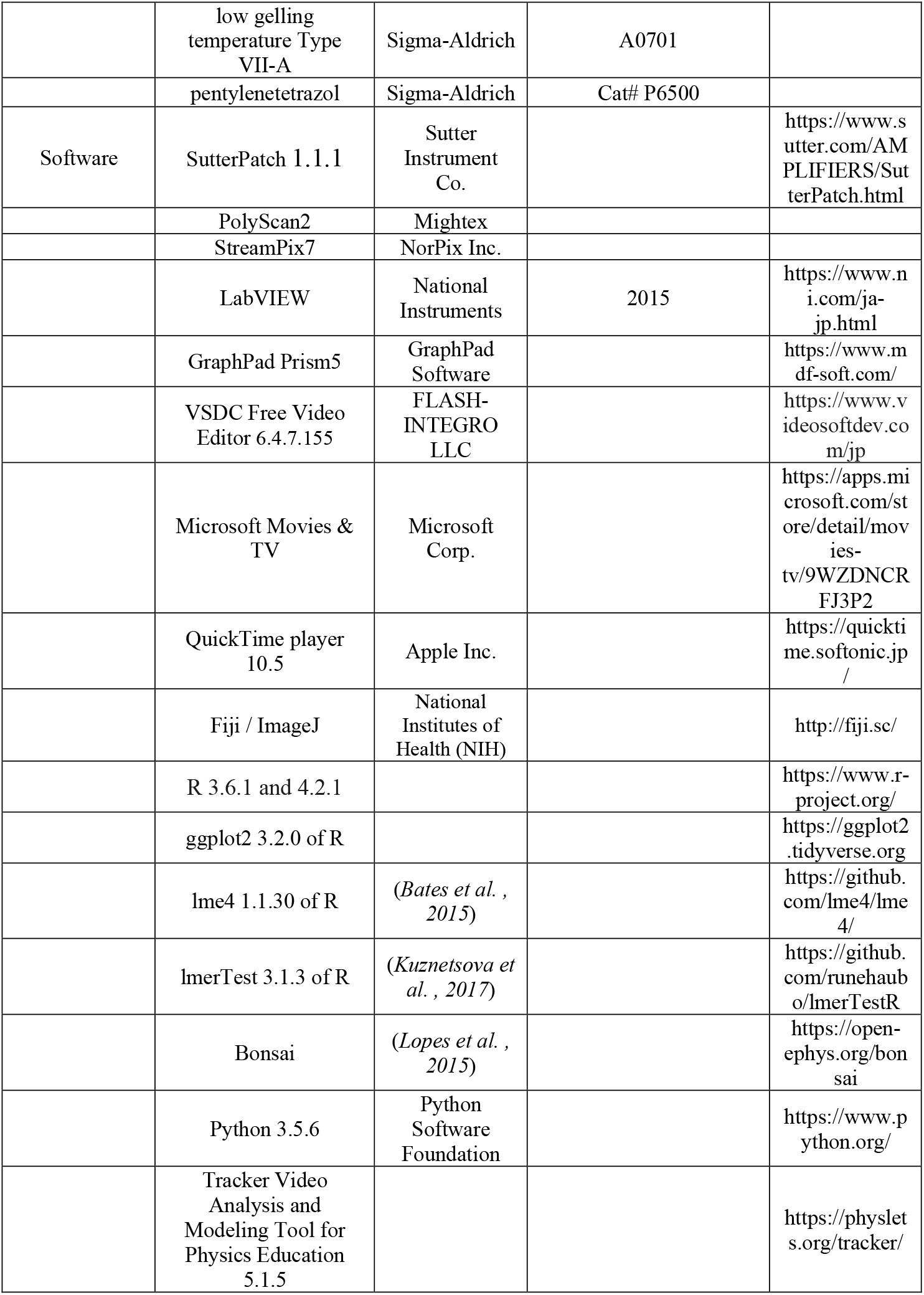

### Ethics statement

The animal experiments in this study were approved by the Nagoya University Animal Experiment Committee and were conducted in accordance with the Regulation on Animal Experiments from Nagoya University.

### Cell culture

The electrophysiological assays of channelrhodopsins were performed on ND7/23 cells, which are hybrid cell lines derived from neonatal rat dorsal root ganglia neurons fused with mouse neuroblastoma (*Wood et al., 1990*). ND7/23 cells were grown on a collagen-coated coverslip in Dulbecco’s modified Eagle’s medium (FUJIFILM Wako Pure Chemical Corp., Osaka, Japan) supplemented with 2.5 μM all-*trans* retinal, 5% fetal bovine serum under a 5% CO2 atmosphere at 37°C. The expression plasmids were constructed based on pCS2+ (see the section of Zebrafish) and were transiently transfected by using FuGENE HD (Promega, Madison, WI, USA) according to the manufacturer’s instructions. Electrophysiological recordings were then conducted 16-36 h after transfection. Successfully transfected cells were identified by EYFP, CFP, or mCherry fluorescence under a microscope prior to measurements.

### Electrophysiology

All experiments were carried out at room temperature (25 ± 2°C). Photocurrents were recorded using an amplifier IPA (Sutter Instrument Co., Novato, CA, USA) under a whole-cell patch clamp configuration. Data were filtered at 5 kHz and sampled at 10 kHz and stored in a computer (Sutter Instrument Co.). The standard internal pipette solution for whole-cell voltage-clamp contained (in mM) 125 K-gluconate, 10 NaCl, 0.2 EGTA, 10 HEPES, 1 MgCl_2_, 3 MgATP, 0.3 Na2GTP, 10 Na2-phosphocreatine, 0.1 Leupeptin, adjusted to pH 7.4 with KOH. The standard extracellular solution contained (in mM) 138 NaCl, 3 KCl, 10 HEPES, 4 NaOH, 2 CaCl_2_, 1 MgCl_2_, 11 glucose, adjusted to pH 7.4 with KOH. Time constants were determined by a single exponential fit, unless noted otherwise.

### Optics for cultured cells

For the whole-cell patch clamp, irradiation at 385, 423, 469, 511, 555, 590 or 631 nm was carried out using Colibri7 (Carl Zeiss, Oberkochen, Germany) controlled by computer software (SutterPatch Software version 1.1.1, Sutter Instrument Co.). Light power was directly measured under an objective lens of the microscope by a visible light-sensing thermopile (MIR-100Q, SSC Inc., Mie, Japan).

### Zebrafish

All transgenic zebrafish lines in this study were generated using the *mitfa^w2/w2^* mutant (also known as *nacre*) line, which lacks melanophores (*Lister et al., 1999*). To construct expression plasmids for *Gt*CCR4, the cDNA of a fusion protein *Gt*CCR4-3.0-EYFP or the open reading frame (ORF) of *Gt*CCR4 with a MycTag (MT) tag sequence, the 2A peptide sequence (P2A) from porcine teschovirus (PTV-1) (*Provost et al., 2007, Tanabe et al., 2010*), and TagCFP (Everon) (*Gt*CCR4-MT-P2A-TagCFP) were subcloned to pCS2+. *Gt*CCR4-3.0-EYFP, which contains a membrane-trafficking signal and the ER-export signal (3.0) from the Kir2.1 potassium channel (*Gradinaru et al., 2010, Hoque et al., 2016*), was constructed according to a previously described procedure (*Hoque et al., 2016*). To construct *Kn*ChR expression plasmids, a carboxy terminal-truncated version (amino acids 1-272 were used), fused with the membrane- and ER-export signals and EYFP, was used (*Tashiro et al., 2021*). The *Kn*ChR-3.0-EYFP, *Cr*ChR2[T159C]-mCherry, *Gt*ACR1-EGFP, *Be*GC1-EGFP, *OaPAC*, and *b*PAC-MT-T2A (2A sequence from *Thosea asigna* virus) DNA fragments were isolated by PCR from phKnChhR-272-3.0-eYFP (*Tashiro et al., 2021*), pmCherry ChR2 T159C (*Berndt et al., 2011*), pFUGW-h*Gt*ACR1-EYFP (*Govorunova et al., 2015*) (a gift from John Spudich [RRID:ADDgene_67795]), peGFP-N1-BeGC1 (*Matsubara et al., 2021*), pCold His *Oa*PAC (*Ohki et al., 2016*) (a gift from Sam-Yong Park), and pAAV hSyn1 bPAC cMyc T2A tDimer (a gift from Thomas Oertner [RRID:Addgene_85397]), respectively, and were subcloned to pCS2+ (for *Oa*PAC, Flag-P2A-TagCFP was attached at the carboxyterminal). pENTR L1-R5 entry vectors containing five repeats of the upstream activation sequence (UAS) and the *hsp70l* promoter (*Muto et al., 2017*), and pENTR L5-L2 vectors containing the ORF of the optogenetic tools and the polyadenylation site of SV40 (SV40pAS) from the pCS2+ plasmids were generated by the BP reaction of the Gateway system. The UAS-hsp70l promoter and optogenetic tool expression cassettes were subcloned to the Tol2 donor vector pBleeding Heart (pBH)-R1-R2 (*Dohaku et al., 2019*), which was modified from pBH-R4-R2 and contains the mCherry cDNA and SV40 pAS under the *myosin, light chain 7, regulatory* (*myl7*) promoter (*van Ham et al., 2010*) by the LR reaction of the Gateway. To make a Tol2 plasmid expressing the cAMP fluorescent indicator Flamindo 2 in all postmitotic neurons, the *elavl3* promoter (*Park et al., 2000*), Flamindo 2 cDNA (*Odaka et al., 2014*), and SV40pAS were subcloned to pT2ALR-Dest (pT2ALR-elav3-Flamindo2). To make Tg fish, 25 pg of the Tol2 plasmids and 25 pg of transposase capped and polyadenylated RNA were injected into one-cellstage embryos. The Tg fish that expressed optogenetic tools in a Gal4-dependent manner are referred as to *Tg(UAS:opto-tool). Tg(UAS:opto-tool*) fish were crossed with *TgBAC(vsx2:GAL4FF*) (*Kimura et al., 2013*), *Tg(myl7:GAL4FF*), or *Tg(elavl3:GAL4-VP16) (Kimura et al., 2013*) to express tools in the hindbrain reticulospinal V2a neurons, heart, and all postmitotic neurons, respectively. For Ca^2+^ imaging and cAMP monitoring, *Tg(5xUAS-hsp70l:GCaMP6s) (Muto et al., 2017*) and *Tg(elavl3:Flamindo2*) were used. Adult zebrafish were raised at 28.5°C with a 14 h light and 10 h dark cycle. Individual larvae used for behavioral experiments were kept in the dark except for the fluorescence observation and light exposure experiments.

### Immunostaining

For immunostaining, anti-Flag antibody (1:500, mouse, Sigma-Aldrich, St. Louis, MO, USA, Cat# F3165, RRID:AB_259529), anti-Myc Tag (MT) antibody (1:500, mouse, Santa Cruz Biotechnology, Dallas, TX, USA, Cat# sc-40, RRID:AB_627268), anti-GFP (1:500, rat, Nacalai Tesque, Inc., Kyoto, Japan, Code: 04404-84, RRID:AB_10013361), and anti-DsRed (1:500, rabbit, Takara Bio, Shiga, Japan, Cat# 632496; RRID:AB_10013483) were used as primary antibodies. CF488A anti-mouse IgG (1:500, H + L, Biotium Inc., Fremont, CA, USA, Cat# 20018; RRID:AB_10557263), CF488A anti-rat IgG (1:500, H + L, Biotium, Inc., Cat# 20023; RRID:AB_10854384) and CF568 anti-rabbit IgG (1:500, H + L, Biotium Inc., Cat# 20103; RRID:AB_10558012) were used as secondary antibodies. The detailed method for immunostaining is described in the accompanying paper. Images were acquired using a confocal laser inverted microscope LSM700 (Carl Zeiss, Oberkochen, Germany). To detect weak fluorescent signals, laser power was increased, but when the power was increased by a factor of 2 or more, it was noted in the figure legend (Figure 6A).

### Locomotion assay

The expression of optogenetic tools in 3-dpf larvae was determined by the expression of fluorescent marker in reticulospinal V2a neurons. Sibling fish that did not express the fluorescent marker were used as control fish. The detailed method is described in the accompanying paper. Briefly, after larvae were anesthetized with tricaine methansulfonate (Nacalai Tesque, Inc., Kyoto, Japan, Cat# 01916-32) and embedded in 2.5% agarose (low gelling temperature Type VII-A A0701, Sigma-Aldrich), the tail was set free by cutting the agarose around it. This agarose was placed in a 90-mm Petri dish filled with rearing water and kept for 20 min to recover from anesthesia. Light stimulation was performed using a patterned LED illuminator system LEOPARD (OPTO-LINE, Inc., Saitama, Japan) and the control software PolyScan2 (Mightex, Toronto, Canada) was used. The irradiation intensity and area were 0.4 mW/mm^2^ and 0.3 mm × 0.34 mm. Tail movements were captured by an infrared CMOS camera (67 fps, GZL-CL-41C6M-C, Point Grey, Canada) mounted under the stage and StreamPix7 software (NorPix Inc., Montreal, Canada) and analyzed by Tracker Video Analysis and Modeling Tool for Physics Education version 5.1.5. The timing of tail motion capture and light irradiation to the reticulospinal V2a neurons was controlled by a USB DAQ device (USB-6008, National Instruments, Austin, TX, USA) and the programming software LabVIEW (2015, National Instruments). The stimulation was repeated six times every 20 min, 100 ms (channelrhodopsins) or 500 ms (adenylyl cyclase) each time, with a minimum of eight individuals for each strain. Trials in which swimming behavior was induced within 8 s after light stimulation were defined as induced trials. The percentage of induced trials was defined as locomotion rate, excluding trials in which swimming behavior was elicited before light stimulation. The time from the start of light irradiation to the first tail movement was defined as latency, and the time from the start of the first tail movement to the end of that movement was defined as duration. The maximum distance the tail moved from the midline divided by the body length was defined as strength. To examine the tools’ ability to inhibit locomotion, 4-dpf Tg larvae were pretreated with 15 mM pentylenetetrazol (Sigma-Aldrich, Cat# P6500) and spontaneous tail movements were induced by white LED light (peak 640 nm; Kingbright Electronic Co., Ltd., Taipei Hsien, Taiwan) powered by a DC power supply (E3631A; Agilent Technologies, Santa Clara, CA, USA) for 5 s. After 500 ms from the onset of the tail movement, the hindbrain reticulospinal V2a were stimulated with the patterned LED illuminator. Trials in which swimming behavior stopped within 1 s after light stimulation were defined as locomotioninhibition trials. The percentage of locomotion-inhibition trials was calculated and indicated in Table 1. Graphs were created with GraphPad Prism5 software (GraphPad Software, San Diego, CA, USA). We used VSDC Free Video Editor version 6.4.7.155 (FLASH-INTEGRO LLC, Moscow, Russia) and Microsoft Movies & TV (Microsoft Corp., Redmond, WA, USA) to make all movies.

### Heartbeat experiments

4-dpf larvae carrying an expressed fluorescent marker in the heart were used for the experiments. Sibling fish that did not express the marker were used as control fish. Four larvae were used for each line. The detailed method is described in the accompanying paper. Briefly, after larvae were quickly anesthetized with about 0.2% tricaine methanesulfonate and embedded in agarose, they were placed in a 90-mm Petri dish filled with water and kept for 20 min to recover from anesthesia. Irradiation intensity was adjusted to 0.5 mW/mm^2^. The area of irradiation was 0.17 mm × 0.25 mm, including the entire heart. The heart area in Tg fish was irradiated for 100 ms (channelrhodopsins) or 5 s (*b*PAC-MT) with light wavelengths that had the closest values to the maximum absorption wavelength of each optogenetic tool. The HBs of larvae were captured by an infrared CMOS camera (67 fps) and recorded with StreamPix7 software as described above. The irradiation trial was repeated six times every 3 min (for *Gt*CCR4-3.0-EYFP and *Gt*ACR1-EYFP) or 10 min (for *Kn*ChR-3.0-EYFP, *Cr*ChR2[T159C]-mCherry, and *b*PAC-MT) for one fish and a total of four larvae were analyzed for each strain. The video recordings of the HBs were observed using the QuickTime player 10.5 (Apple Inc., Cupertino, CA, USA). After opening videos with Fiji/ImageJ (National Institutes of Health, Bethesda, MD, USA), the entire heart was set as the region of interest (ROI), the luminosity (AU: arbitrary units) data in the ROI was used to create graphs of HBs using ggplot2 version 3.2.0 in R. The relative HB frequency was calculated by Bonsai (*Lopes et al., 2015*) and Python version 3.5.6 (Python Software Foundation, Wilmington, DE, USA). Graphs of the average of relative HB frequency were created by ggplot2 of R. The latency to cardiac arrest and the time to first resumption of HB were also measured. Graphs were created with GraphPad Prism5 software. All movies were created with VSDC Free Video Editor. Simple HB experiments were also performed using a light source equipped with an MZ16 FA microscope and GFP (460-500 nm), YFP (490-510 nm), and DSR filters (530-560 nm, Leica, Wetzlar, Germany), as indicated in Table 1.

### Ca^2+^ imaging

4-dpf Tg fish expressing *Kn*ChR or *Gt*ACR1, and GCaMP6s in cardiomyocytes were used. Tg fish expressing only GCaMP6s were used as controls. The larvae anesthetized with tricaine methanesulfonate were embedded in 3% agarose (low gelling temperature Type VII-A, Sigma-Aldrich) in 1/10 Evans solution, placed in a 90-mm Petri dish filled with water, and left on the microscope stage for 10 min. A 130 W light source (U-HGLGPS, Olympus, Tokyo, Japan) with the fluorescence detection filter (excitation 470-495 nm, emission 510-550 nm, U-MNIBA3, Olympus) was used to observe the fluorescence of GCaMP6s. A CCD camera (ORCA-R2, Hamamatsu Photonics, Shizuoka, Japan) located on the microscope was used to capture the GCaMP6s fluorescence images at 9 fps. After image acquisition, the entire heart area was manually set as the ROI using Fiji/ImageJ, and fluorescence intensity was measured. Trials were repeated three times every 10 min. The relative change in ΔF/F was calculated by dividing the fluorescence intensity in each frame by the fluorescence intensity at the start of light exposure.

### cAMP live imaging

3-dpf larvae expressing an optogenetic tool and the cAMP indicator Flamindo2 in postmitotic neurons were used. Sibling larvae that did not express the optogenetic tool were used as controls. The larvae that were quickly anesthetized with 0.04% tricaine methanesulfonate were embedded in 3% agarose, placed in a 90-mm Petri dish filled with rearing water, and left on the microscope stage for 20 min. A 130 W light source (U-HGLGPS, Olympus) with a fluorescence detection filter (excitation 470-495 nm, emission 510-550 nm) was used for observation. The fluorescence images were captured by a CCD camera (ORCA-R2, Hamamatsu Photonics) at 9 fps. After image acquisition, the entire optic tectum area was set as a ROI using ImageJ and fluorescence intensity was measured. ΔF/F was calculated.

### Statistical analysis

Data were analyzed using R software package (versions 3.6.1 and 4.2.1). Statistical tests were applied as indicated in figure legends. All data in the text and figures are expressed as the mean ± standard error. A linear mixed-effects model was applied using R package ‘lme4’ version 1.1.30 and ‘lmerTest’ version 3.1.3 (*Bates et al., 2015, Kuznetsova et al., 2017*). The *p*-values of this model were computed with a *t* test based on the degrees of freedom with Scatterthwaite’s method, including time, strain as the fixed effect, and individuals as the random effect.

## Acknowledgements

We thank Shin-ichi Higashijima, Koichi Kawakami, and the National Bioresource Project for providing transgenic fish; John Spudich, Sam-Yong Park, and Thomas Oertner for providing plasmid DNAs; Tamiko Itoh for managing fish mating and care; Ryosuke Takeuchi for helping us analyze heartbeat experiments. We also thank the members of the Kandori and Hibi laboratories for helpful discussion.

## Figure legends

**Figure S1.**
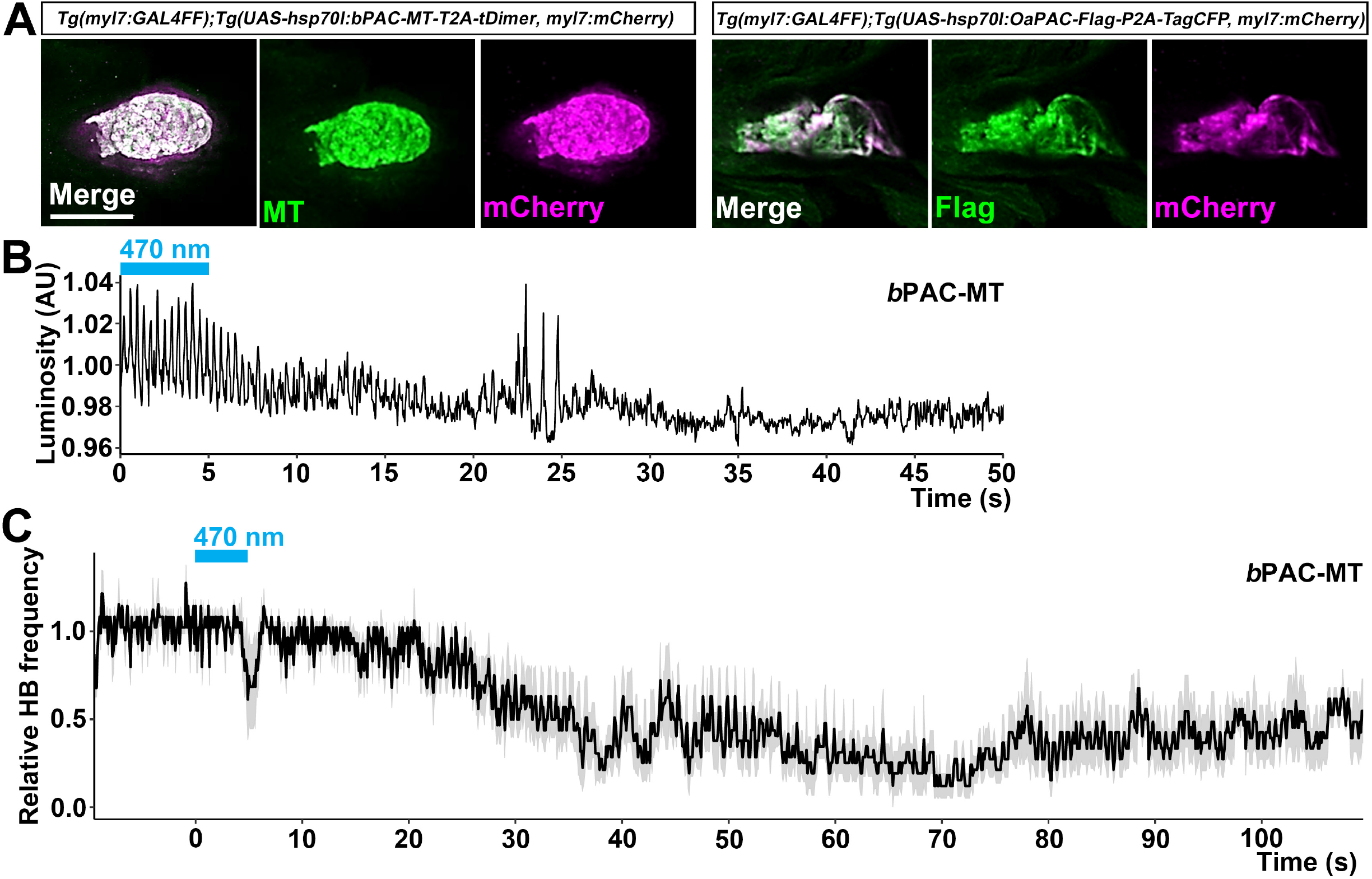
Optogenetic control of the heart by *b*PAC or *Oa*PAC. (A) Expression of *b*PAC-MT or *Oa*PAC-Flag in cardiomyocytes. 4-dpf larvae expressing *b*PAC-MT or *Oa*PAC-Flag were fixed and stained with anti-Myc or Flag (green), and anti-DsRed (mCherry, magenta) antibodies. (B, C) HBs monitored by luminosity (AU), changes (B), and relative HB frequency (C) of *b*PAC-expressing larvae. The heart area of Tg larvae expressing *b*PAC was irradiated with light (0.5 mW/mm^2^) of 470 nm for 5 s at the indicated periods. Similar results were obtained from four Tg larvae. A typical example from one larva is shown in (B), and average HB frequency of the 1st or 2nd trial showing a typical pattern for the four larvae is shown in (C). The larvae showed induced bradycardia in the 3rd through 6th trials. Scale bar = 100 μm in (A).

## Movies

**Movie 1.** Tail movements in a larva expressing *Cr*ChR2[T159C]-mCherry in reticulospinal V2a neurons.

The hindbrain in a 3-dpf *Tg(vsx2:GAL4FF); Tg(UAS-hsp70l:CrChR2[T159C]-mCherry, myl7:mCherry*) larva was stimulated with 470 nm light for 100 ms. The timing of light stimulation is indicated by a blue circle.

**Movie 2.** Tail movements in a larva expressing *Gt*CCR4-3.0-EYFP in reticulospinal V2a neurons.

The hindbrain in a 3-dpf *Tg(vsx2:GAL4FF); Tg(UAS-hsp70l:GtCCR4-3.0-EYFP, myl7:mCherry*) larva was stimulated with 520 nm light for 100 ms. The timing of light stimulation is indicated by a green circle.

**Movie 3.** Tail movements in a larva expressing *Gt*CCR4-MT-P2A-TagCFP in reticulospinal V2a neurons.

The hindbrain in a 3-dpf *Tg(vsx2:GAL4FF);Tg(UAS-hsp70l:GtCCR4-MT-P2A-TagCFP, myl7:mCherry*) larva was stimulated with 520 nm light for 100 ms. The timing of light stimulation is indicated by a green circle.

**Movie 4.** Tail movements in a larva expressing *Kn*ChR-3.0-EYFP in reticulospinal V2a neurons.

The hindbrain in a 3-dpf *Tg(vsx2:GAL4FF);Tg(UAS-hsp70l:KnChR-3.0-EYFP, myl7:mCherry*) larva was stimulated with 470 nm light for 100 ms. The timing of light stimulation is indicated by a blue circle.

**Movie 5.** Heart movements in a larva expressing *Gt*CCR4-3.0-EYFP in cardiomyocytes. The heart area of *Tg(myl7:GAL4FF);Tg(UAS-hsp70l:GtCCR4-3.0-EYFP, myl7:mCherry*) was stimulated with 520 nm light for 100 ms. The timing of light stimulation is indicated by a green circle.

**Movie 6.** Heart movements in a larva expressing *Kn*ChR-3.0-EYFP in cardiomyocytes.

The heart area of *Tg(myl7:GAL4FF);Tg(UAS-hsp70l:KnChR-3.0-EYFP, myl7:mCherry*)was stimulated with 470 nm light for 100 ms. The timing of light stimulation is indicated by a blue circle.

**Movie 7.** Heart movements in a larva expressing *Cr*ChR2[T159C]-mCherry in cardiomyocytes.

The heart area of *Tg(myl7:GAL4FF);Tg(UAS-hsp70l:CrChR2[T159C]-mCherry, myl7:mCherry*) was stimulated with 470 nm light for 100 ms. The timing of light stimulation is indicated by a blue circle.

**Movie 8.** Heart movements in a larva expressing *Gt*ACR1-EYFP in cardiomyocytes.

The heart area of *Tg(myl7:GAL4FF);Tg(UAS-hsp70l:GtACR1-EYFP, myl7:mCherry*)was stimulated with 520 nm light for 100 ms. The timing of light stimulation is indicated by a green circle.

**Movie 9.** Ca^2+^ imaging in the heart of a larva expressing *Kn*ChR-3.0-EYFP and GCaMP6s in cardiomyocytes.

The heart area of *Tg(myl7:GAL4FF);Tg(UAS-hsp70l:KnChR3.0-EYFP, myl7:mCherry);Tg(UAS-hsp70l:GCaMP6s*) was stimulated with a fluorescence detection filter (excitation 470-495 nm, emission 510-550 nm). GCaMP6s fluorescence was monitored with the same filter set. The fluorescence of GCaMP6s gradually increased.

**Movie 10.** Ca^2+^ imaging in the heart of a larva expressing *Gt*ACR1-EYFP and GCaMP6s in cardiomyocytes.

The heart area of *Tg(myl7:GAL4FF);Tg(UAS-hsp70l:GtACR1-EYFP, myl7:mCherry);Tg(UAS-hsp70l:GCaMP6s*) was stimulated with a fluorescence detection filter (excitation 470-495 nm, emission 510-550 nm). GCaMP6s’ fluorescence was monitored with the same filter set. The fluorescence of GCaMP6s gradually decreased.

**Movie 11.** Tail movements in a larva expressing *Be*GC1-EGFP in reticulospinal V2a neurons.

The hindbrain in a 3-dpf *Tg(vsx2:GAL4FF);Tg(UAS-hsp70l:BeGC1-EGFP, myl7:mCherry*) larva was stimulated with 520 nm light for 500 ms. The timing of light stimulation is indicated by a green circle.

**Movie 12.** Tail movements in a larva expressing *b*PAC-MT in reticulospinal V2a neurons. The hindbrain in a 3-dpf *Tg(vsx2:GAL4FF);Tg(UAS-hsp70l:bPAC-MT-T2A-tDimer, myl7:mCherry*) larva was stimulated with 470 nm light for 500 ms. The timing of light stimulation is indicated by a blue circle.

**Movie 13.** Tail movements in a larva expressing *Oa*PAC-Flag in reticulospinal V2a neurons.

The hindbrain in a 3-dpf *Tg(vsx2:GAL4FF);Tg(UAS-hsp70l:OaPAC-Flag-P2A-TagCFP, myl7:mCherry*) larva was stimulated with 470 nm light for 500 ms. The timing of light stimulation is indicated by a blue circle.

**Movie 14.** Heart movements in a larva expressing *b*PAC-MT in cardiomyocytes.

The heart area of *Tg(myl7:GAL4FF);Tg(UAS-hsp70l:bPAC-MT-T2A-tDimer, myl7:mCherry*) was stimulated with 470 nm light for 5 s. The timing of light stimulation is indicated by a blue circle.

